# Inter-Individual and Inter-Strain Differences in Cognitive and Social Abilities of Dark Agouti and Wistar Han Rats

**DOI:** 10.1101/566877

**Authors:** Lucille Alonso, Polina Peeva, Arnau Ramos-Prats, Natalia Alenina, York Winter, Marion Rivalan

## Abstract

**Background:** Healthy animals showing extreme behaviours spontaneously that resemble human psychiatric symptoms are relevant models to study the natural psychobiological processes of maladapted behaviours. Healthy poor decision makers (PDMs) identified using a Rat Gambling Task, co-express a combination of cognitive and reward-based characteristics similar to symptoms observed in human patients with impulse-control disorders. The main goals of this study were to 1) confirm the existence of PDMs and their unique behavioural phenotypes in the Dark Agouti (DA) and Wistar Han (WH), 2) to extend the behavioural profile of the PDMs to probability-based decision-making and social behaviours and 3) to discuss how the key traits of each strain could be relevant for biomedical research.

**Methods:** We compared cognitive abilities, natural behaviours and physiological responses in DA and WH rats using several tests. We analysed the results at the strain and the individual level.

**Results:** Previous findings in WH rats were reproduced and could be generalized to DA. Each PDM of either strain displayed a similar, naturally occurring, combination of behavioural traits, including possibly higher social rank, but no deficits in probability-based decision-making. A Random forest analysis revealed interesting discriminating traits between WH and DA.

**Conclusion:** The reproducibility and conservation of the socio-cognitive and behavioural phenotypes of GDM (good decision maker) and PDM individuals in the two genetically different strains of WH and DA support a good translational validity of these phenotypes. Both DA and WH rat strains present large phenotypic variations in behaviour pertinent for the study of the underlying mechanisms of poor decision making and associated disorders.

## 1. Introduction

Inter-individual variability in behaviour is a natural phenomenon that applies to all behavioural dimensions. In the laboratory, however, these phenotypic variations are often perceived as inconvenient and are usually masked by averaging of the data. Considering the spectrum nature of brain disorders, most psychiatric symptoms can be conceptualized as extreme manifestations of different behavioural traits [1]. Thus, the identification of animals spontaneously exhibiting extreme behaviours that resemble human psychiatric symptoms offers the opportunity to study the natural psychobiological processes underlying maladapted behaviours [2,3].

Utilizing this dimensional approach to the analysis of the Rat Gambling Task (RGT), a rat version of the human Iowa Gambling task, we and others have consistently identified three types of decision makers spontaneously existent in healthy groups of Wistar Han (WH) and Sprague Dawley rats [4–8]. Whereas the majority of rats develop a strong preference for the most advantageous options in the RGT (good decision makers [GDMs]), a smaller group prefer the least advantageous options (poor decision makers [PDMs]) and some show no clear preference (intermediate phenotype [INT]) [6].

Compared to GDMs, healthy PDMs were found to co-express several cognitive impairments and reward-based deficits similar to symptoms observed in human patients with substance abuse disorder, pathological gambling disorder, attention-deficit hyperactivity-disorder (ADHD) or suicidal behaviour [6,8,9]. Healthy PDMs were more prone to take risks in potentially dangerous environments, showed higher motivation to obtain a reward and greater anticipatory (motor) impulsive responses, were more inflexible and chose less advantageously in the RGT due to their over-valuation of the high-reward/high-risk options compared with GDMs [8]. Their social abilities and spontaneous level of activity (e.g. arousal) are, however, still unknown [10,11]. At the biological level, PDMs also presented a particular profile compared to GDMs. PDMs showed different use of distinct regions of the prefrontal cortex (PFC) to solve the RGT [7], a decreased c-Fos activation in the PFC-subcortical network normally used by the GDMs [5] and an opposite pattern of serotonin turnover compared to GDMs, with higher turnover rate in the PFC (i.e. infralimbic cortex) but lower turnover rate in subcortical areas (i.e. basolateral amygdala) [5].

Among other candidates, the serotonergic system appears to be a promising pathway that could be responsible for the co-expression of the traits constitutive of the PDM psychobiological profile. Indeed, serotonin plays a critical role in executive functioning (decision making, impulse control, flexibility, attention), mood control, sociality and emotional state [9,12–19], and is a privileged therapeutic target for treating pathologies associated with poor decision making such as substance abuse, ADHD, suicidal behaviour, impulsive control disorders (i.e., eating disorders, gambling), psychopathy and other aggression related disorders [20–22]. Although more than one behavioural domain was rarely tested in the same individual, other studies have reported equivalent deleterious effects of the dietary, genetic or pharmacological reductions of central serotonin function on group (*vs.* inter-individual) performance in decision making [23,24], motor impulsivity [25] and cognitive inflexibility [26], but also in social recognition [27], aggression [28] and social hierarchy [29,30].

In order to evaluate the functional role of the serotonergic system in the expression of the vulnerable behavioural profile in rats, we plan to use an animal model of congenital central serotonin depletion [31]. The background strain of this newly created rat line is the Dark Agouti (DA) strain. However, historically, DA rats have been mainly used in physiological studies, and have only rarely been tested for their cognitive abilities [32] and never for their social skills. We also wanted to confirm that this inbred strain of rats naturally displayed comparable behavioural phenotypic variability to WH [33].

Therefore, the goal of this study was to evaluate the conservation of the GDM and PDM profiles between the WH and DA strains by establishing the bio-behavioural profile of the DA rats, examining the same behavioural traits naturally exhibited by the WH rats. We also used this opportunity to test the reproducibility of previous results obtained from a different laboratory with the WH strain, and to extend the behavioural profile of the PDMs to serotonin-sensitive tasks such as probability based decision making and social behaviours. We compared cognitive abilities, natural behaviours and physiological responses in DA and WH rats using several tests. These tests included the RGT, the reversed-RGT, the Delay discounting task (DDT), the Probability discounting task (PDT), the Fixed-interval and Extinction schedule of reinforcement (FI-EXT), a semi-automated version of the Visible Burrow System (VBS), the Social Recognition test (SRt) and the Elevated Plus maze (EPM). The results were analysed at both the group (strain) and individual (within strain) levels. Finally, by performing a random forest analysis, we were able to highlight key traits to discriminate one strain from the other and discuss the relevance of using each strain in different types of studies.

## 2. Material and Methods

### 2.1. Animals

In this study, we used 42 male WH rats (Charles River, Germany) and 42 male DA rats (Max Delbrück Center for Molecular Medicine, Berlin). They arrived at our animal facility at between six and nine weeks of age. Rats of the same strain were housed in pairs in standard rat cages (Eurostandard Type IV, 38 cm x 59 cm) in two temperature-controlled rooms (22°C and 50% humidity) with inverted 12-hour light cycles (lights on at 20:00 in room 1 or 01:00 in room 2). The two different light cycles allowed us to maximize the use of four operant cages with two groups of 12 animals tested either in the morning or in the afternoon (i.e. 24 animals per day). To habituate the animals to their new environment, they were left undisturbed for at least a week after arrival. Thereafter, they were handled daily by the experimenter. Two weeks before the beginning of the training phase, rats were marked subcutaneously with a radio-frequency identification (RFID) chip (glass transponder 3 mm, Euro I.D.) under short isoflurane anaesthesia. Rats were between 9 and 12 weeks of age when first trained in the operant cages. Rats had *ad libitum* access to food and water. During operant training and testing, rats were maintained at 95% of free-feeding weight by food restriction. One DA rat was excluded from the RGT and reversed-RGT analysis since it did not show sampling behaviour at the start of the test and a strong side bias over the entire duration of the tests. One DA extreme outlier (< mean – 2*SD) was excluded from the weight analysis after VBS housing.

### 2.2. Ethics

All procedures followed national regulations in accordance with the European Communities Council Directive 2010/63/EU. The protocols were approved by the local animal care and use committee and run under the supervision of the animal welfare officer of the animal facility of the Charité University Medicine.

### 2.3. Behavioural tests

Training and testing started 1 h after the beginning of the dark phase. Animals were habituated to the experimental room conditions for 30 min. The order of the tests and inter-test pauses was chosen to minimize any interference of one test on another (Fig. 1A). One group of 12 WH rats performed the DDT before the VBS housing. Not all animals underwent all tests (as can be seen from the different numbers of animals in the figures).

**Figure 1.**
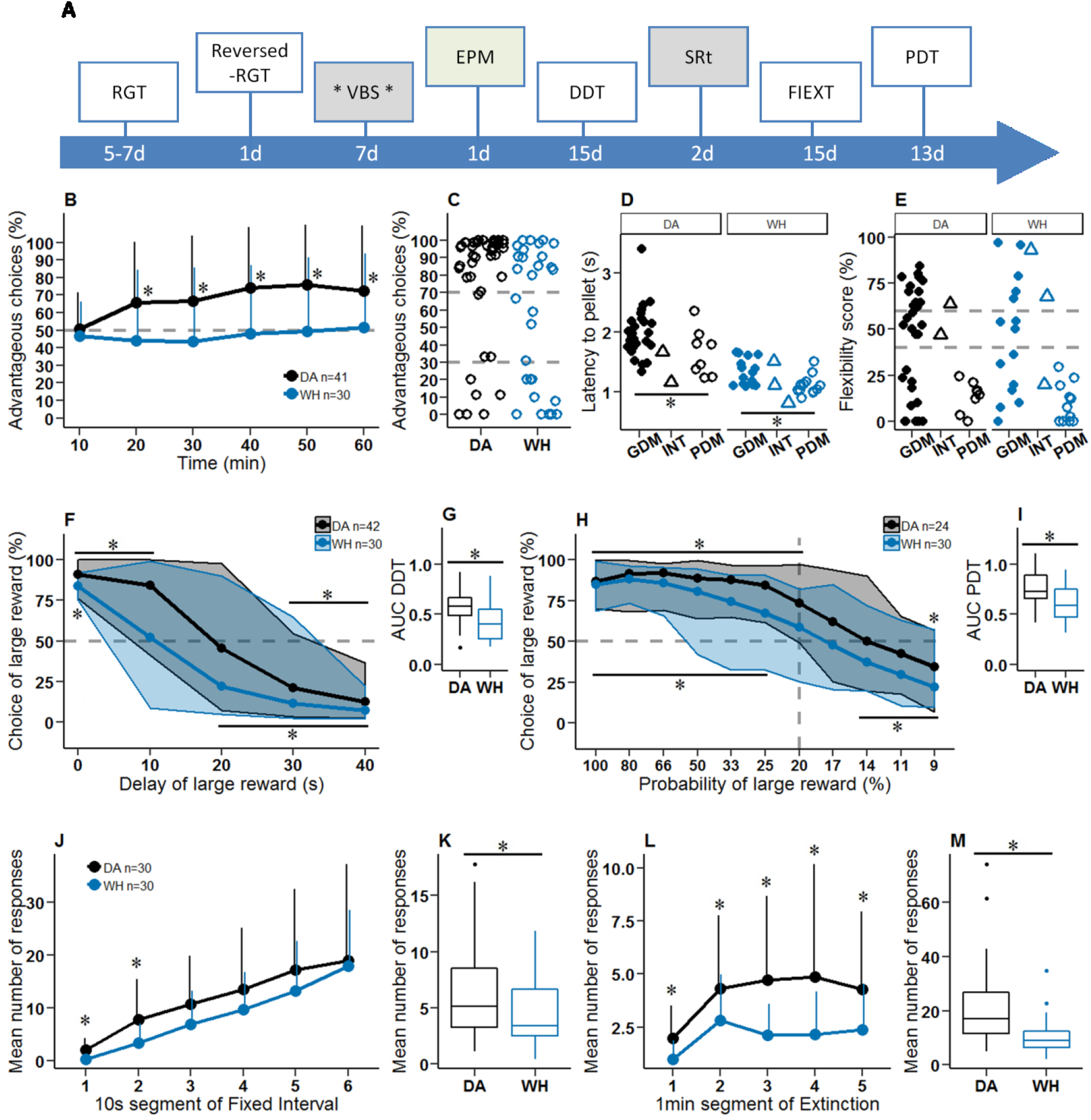
Order and duration of testing and cognitive abilities of Dark Agouti (DA) and Wistar Han (WH) rats in the RGT, reversed-RGT, DDT, PDT and FIEXT. **A** Order and duration of testing. RGT: Rat gambling task. VBS: Visible burrow system, with faeces collection (asterisks) before and after VBS housing. EPM: Elevated plus maze. DDT: Delay discounting task. SRt: Social recognition test. PDT: Probability discounting task. Cognitive tasks are in white and social tasks are in grey. d: day. **B** Advantageous choices in the RGT. Data are mean + SD, one sample t-test *vs.* 50%. **C** Individual (mean) scores during the last 20 min of the RGT. The dashed line at 70% and 30% of advantageous choices visually separates good decision makers (GDMs), intermediates (INTs) and poor decision makers (PDMs). **D** Motivation for the reward in the RGT, with filled circles representing GDMs, triangles representing INTs and empty circles representing PDMs; Wilcoxon rank sum test, GDM *vs.* PDM. **E** Flexibility scores in the reversed-RGT. **F** Choice of the large reward option as a function of the delay of reward delivery. Lines indicate the medians, and areas shaded in grey (DA) or blue (WH) indicate the 5^th^ to 95^th^ percentiles. The dashed line indicates the 50% chance level. The asterisk denotes significant difference (Wilcoxon sign test) from 50% choice for DA (*above curve) and WH (*below curve). **G** Area under the curve for the DDT; Wilcoxon rank sum test, DA *vs.* WH. **H** Choice of the large reward option as a function of the probability of reward applied. Lines indicate the median, and areas shaded in grey (DA) or blue (WH) indicate the 5^th^ to 95^th^ percentiles. The vertical dashed line shows the indifference point (20% chance of receiving 5 pellets). The asterisk shows significant difference (Wilcoxon sign test) from 50% choice for DA (*above curve) and WH (*below curve). **I** Area under the curve for the PDT. Wilcoxon rank sum test, DA *vs.* WH. **J** Mean number of nose pokes during the 1 min FI expressed for the 10 s segments + SD. **K** Mean number of nose pokes during all 1 min FI (last 4 days). **L** Mean number of nose pokes during the 5 min EXT expressed for the 1 min segments + SD. Wilcoxon rank sum test, DA *vs.* WH. **M** Mean number of nose pokes during all EXT (last 4 days). * p < 0.05. DA in black and WH in blue.

#### 2.3.1. Operant system and tests

All operant training and testing was done in four operant cages (Imetronic, Pessac, France) controlled by a computer. The operant cages contained a curved wall on one side equipped with one to four nose-poke holes, depending on the test. On the opposite wall, a food magazine was connected to an outside pellet dispenser. 45 mg sweet pellets (5TUL, TestDiet, USA) were used. A clear partition with a central opening was placed in the middle of the cage, ensuring an equal distance to all nose-poke holes from the central opening.

##### 2.3.1.1. Complex decision making in the RGT

The training and testing procedures have been described previously [6]. The operant cages had four nose-poke holes on the operant wall. Training 1 started with the four nose-poke holes lit; a single nose poke by the rat led to the delivery of one pellet, and the lights in the non-selected holes were then turned off until the food magazine was visited and all holes were lit again. Daily training continued until rats obtained 100 pellets in a 30 min session (cut-off criteria). During Training 2, two consecutive nose pokes at the same hole were required to obtain one pellet; this training continued until rats obtained 100 pellets in a 30 min session. After Training 2 and for all subsequent testing phases, rats always had to make two consecutive nose pokes at the same hole for a valid choice. Training 3 was a single 15 min session in which two pellets were delivered after a choice was made, up to a maximum of 30 pellets. A forced training (Training 4) was given to counter any side preferences developed during the training procedure. This training was given when a rat had chosen the holes of one side of the operant wall in more than 60% of choices during the last session of Training 2. During the first phase of Training 4, only the two nose-poke holes on the non-preferred side were lit, and choosing one of them led to the delivery of one pellet. After the collection of the first 15 pellets, the second phase of Training 4 started with all four holes lit. Choosing one hole from the side preferred in Training 2 was rewarded (with one pellet) in only 20% of the cases, whereas choosing from the other (least-preferred) side was rewarded in 80% of the cases. The cut-off criterion was set at a maximum of 50 pellets or 30 min. This training phase usually took between five and seven days, and the RGT was performed the next day.

During the test, the four nose-poke holes were lit and each hole was associated with an amount of reward and a possible penalty (time-out). Two holes on one side were rewarded with two pellets and associated with unpredictable long time-outs (222 s or 444 s; probability of occurrence 50% and 25%, respectively); over the long term, these options were disadvantageous. Two holes on the other side were rewarded with one pellet and associated with unpredictable short time-outs (6 s or 12 s; probability of occurrence 50% and 25%, respectively); over the long term, these options were advantageous. The theoretical gain of pellets for the advantageous options was five times higher than for the disadvantageous options at the end of the test (i.e., 60 min; [6] see Supplement 1). After a choice, the reward was delivered and the selected hole remained lit until a visit to the magazine or for the duration of the time-out. During this time, all the nose-poke holes were inactive. The cut-off criterion was 250 pellets.

The percentage of advantageous choices during the last 20 min of the RGT was used to identify GDMs and PDMs. GDMs were defined as choosing >70% advantageous options and PDMs as choosing <30% advantageous options. Intermediate animals (INTs) chose between 30% and 70% advantageous options and did not show a steady preference for only one type of option at the end of the test. To visualize progression of preference during the RGT, advantageous choices were plotted for 10 min time intervals. In a previous study, fast and slow GDMs were described based on how rapidly they developed a preference for the advantageous options [5]. Fast GDMs chose >70% advantageous options during the first 20 min of the test, whereas slow GDMs stayed < 70%. The motivation to obtain a reward (reward sensitivity) was indicated by the mean latency to visit the feeder after a choice.

##### 2.3.1.2. Cognitive flexibility in the reversed-RGT

Animals were tested in the reversed-RGT 48 h after performing the RGT [6]. For this test, the contingencies associated with the four holes during the RGT were spatially reversed by switching the sides for the advantageous and disadvantageous options. A test was 60 min (or a cut-off of 250 pellets).

A flexibility score was calculated as the preference for the same preferred options during the reversed-RGT and the RGT, which meant choosing holes at the location of the non-preferred option during the RGT. For INTs and GDMs, the flexibility score was determined from the percent of advantageous choices during the last 20 min. For PDMs, the flexibility score was determined from the percent of disadvantageous choices during the last 20 min.

Flexible rats had flexibility scores > 60%, undecided rats had flexibility scores between 60% and 40%, and inflexible rats had flexibility scores < 40%.

##### 2.3.1.3. Cognitive impulsivity in the DDT

For this task, the two outside nose-poke holes (25 cm apart) were used. The operant cages were otherwise identical to the other tests. During the DDT, one nose-poke hole (NP1) was associated with a small immediate reward (one pellet); the second nose-poke hole (NP5) was associated with a large delayed reward (five pellets). The protocol was adapted from Rivalan et al. [8], in which levers were used instead of nose-poke holes.

During training, the large reward was delivered immediately after the choice (0 s delay), which allowed the rats to develop a preference for NP5. After a choice, the selected hole stayed lit for 1 s. The magazine and house lights were turned on during a 60 s time-out. A session lasted for 30 min or until 100 pellets were delivered. A > 70% preference for the large reward option on two consecutive sessions with ≤ 15% difference was required to start the test. At least three training sessions were performed. During the test, choosing NP5 induced the delivery of the large reward after a fixed delay, and NP5 stayed lit for the duration of the delay. After the delivery of the large reward, the magazine and the house lights were turned on for a time-out (60 s minus the duration of the delay). The delay was fixed for one day, but increased by 10 s from 0 s to 40 s after a stability criterion (≤ 10% variation of choice of the large reward during two consecutive sessions) was met. The test sessions lasted for 60 min or until 100 pellets were delivered. The preference for the large delayed reward was calculated as the mean percentage of NP5 choices during the two stable sessions. Individual area under the curve (AUC) was measured to estimate the cognitive impulsivity. The choices for the large delayed reward were normalized to the choice for the large delayed reward during the training phase (0 s delay) and plotted against the normalized delays on the x-axis (from 0 to 1). The AUC was calculated as the sum of the areas of the trapezoids formed by the individual data points and the x-axis following the formula (x2-x1)[(y1+y2)/2], [34]. The total number of nose pokes during the last training session was used as an index of the activity during this test.

##### 2.3.1.4. Cognitive risk-taking in the PDT

For this task, the two outside nose-poke holes (25 cm apart) were used. The operant cages were otherwise identical to the other tests. During the PDT, one hole (NP1) was associated with a small and certain reward (one pellet) and the second hole (NP5) was associated with a large but uncertain reward (five pellets) [24].

During training, choosing NP5 always delivered the large reward (probability P=100%). This allowed rats to develop a preference for NP5. NP1 always delivered one pellet. The reward was delivered 4 s after a choice was made in one of the nose-poke holes, and the hole stayed lit until pellet collection. The reward delivery was followed by a 15 s time-out during which the magazine light was on. A session lasted 25 min or until 200 pellets were delivered. A ≥ 70% preference for the large reward was required to start the test. At least three training sessions were performed. During the test, the delivery of the large reward was associated with a set probability (P = 80%, 66%, 50%, 33%, 25%, 20%, 17%, 14%, 11%, or 9%). The probability was fixed for one day and decreased every day. A session lasted 25 min or until 200 pellets were delivered. For each individual, the AUC was calculated as in the DDT. The preference for the large reward was normalized to the preference during training and plotted against the probability values expressed as odds, with odds = (1/P)-1 and normalized (x-axis from 0 to 1) [35].

##### 2.3.1.5. Motor impulsivity in the FI-EXT schedule of reinforcement

For this task, only the central nose-poke hole was used. The operant cages were otherwise identical to the other tests. The FI consists of two phases: a fixed time interval during which choices are not rewarded, followed by a phase where a choice can be rewarded [8]. The EXT is a longer, fixed time interval during which no choices are rewarded. Both FI and EXT are conditions that cause frustration in the animal. A session consisted of seven FI of variable duration depending on the session and one EXT of 5 min; this pattern was repeated two times within a single session. The maximum number of pellets was 14 during a single session. FI lasted 30 s for the first four sessions, 1 min for the next four sessions, 2 min for the next three sessions and 1 min for the final four sessions. The final four sessions with a 1 min FI were the actual test. During the FI, the house light was on and the central nose-poke hole was inactive. At the end of the FI, the house light turned off and the central nose-poke was lit and became active; two consecutive nose-pokes induced the delivery of one pellet, the central nose-poke light was turned off and the tray light was lit. A visit to the tray induced the start of the next FI. After seven consecutive FI, the EXT period started, with all lights off and no consequences associated with nose poking. The mean number of nose pokes was measured for each FI and EXT period. We summed nose pokes for 10 s intervals during FI to visualize the anticipatory activity of the rats. Likewise, we summed nose pokes for 1 min intervals during EXT to visualize the perseverative activity. As described earlier [36], the data from the first FI of the session and the first FI after the first EXT were excluded because they deviated from the other intervals.

#### 2.3.2. Social behaviour in the VBS

The VBS consisted of an open area (2000P, 61×43 cm, Tecniplast, Italy) extended to the top by high Forex PVC foam and Plexiglas (Modulor, Germany) walls and connected through two transparent tunnels to a burrow system placed into a second Type IV cage (Supplementary Fig. 5I). The burrow system was made of infrared transparent black plastic and consisted of a large chamber, a small chamber and a tunnel system (25 cm x 53 cm). Throughout the test, the burrow system remained in the dark. Food and water were available in the open area. A grid of 32 RFID detectors (PhenoSys, Berlin, GmbH) was placed below the VBS in order to automatically determine individual animal positions using the program PhenoSoft (PhenoSys Berlin, GmbH). An infrared camera (IP-Camera NC-230WF HD 720p, TriVision Tech, USA) above the VBS recorded a 30 s video every 10 min (CamUniversal, CrazyPixels, Germany). The software PhenoSoft ColonyCage (PhenoSys Berlin, GmbH) was used to identify individuals in the videos. Six rats were housed in the VBS for seven days in a humidity- and temperature-controlled room (temperature 22°C to 24°C, humidity 45% to 50%). The behaviours expressed by the animals were scored on the videos of the last two days of the VBS during the first 4 h of each dark and light phase (100 videos) using a scan sampling method [37]. Four classes of behaviours were scored: affiliative, aggressive, defensive and maintenance (details in Table 1). The behaviours with a median of < 5 occurrences per strain were grouped for the analysis. The body weight of the animals was measured before and after the VBS housing. Although wounds were rarely observed during this study, they were counted and documented at the end of the VBS housing. The activity (distance travelled) and the place preference were extracted using the software PhenoSoft analytics (PhenoSys Berlin, GmbH). The time spent in the open area of the VBS was measured using the data collected from the grid of detectors.

**Table 1.** Ethogram of the behaviours scored during the VBS housing. Based on Burman et al., Rademacher et al., and Whishaw, Ian Q and Kolb Bryan [38–40].

| Category | Behaviour | Definition |
| --- | --- | --- |
| Affiliative | Allogrooming | Gentle grooming of another rat which is not pinned on its back |
| Affiliative | Attending | Orienting the head, ears and possibly the whole body toward another rat |
| Affiliative | Huddle | Lying in contact with another rat |
| Aggressive | Aggressive grooming | Vigorous grooming of another rat while pinning it |
| Aggressive | Attack bite | Sudden bite toward neck and back of another rat |
| Aggressive | Attack jump | Sudden jump toward another rat |
| Aggressive | Following | Rat runs after another one |
| Aggressive | Fight | Rough-and-tumble of two animals |
| Aggressive | Lateral attack | Arched-back posture oriented towards another rat, often including shoving and piloerection |
| Aggressive | Mutual upright posture | Both rats are standing in front of each other with vertical movements of the forepaw |
| Aggressive | Pinning | Being above another rat and maintaining it with the forepaw usually lying on its back |
| Aggressive | Struggle at feeder | Rats are pushing each other to have the place at the feeder |
| Aggressive | Struggle in tunnels | Rats are pushing each other to pass in the tunnel, struggling with the paws. |
| Defensive | Flight | Rapid movement away from another rat |
| Defensive | Freezing | Being immobile or maintaining a specific posture (crouching) |
| Defensive | Lateral defence | Exposing the flank to another rat. |
| Defensive | Supine posture | Lying on the back (exposure of the belly) because of another rat |
| Defensive | Upright defence | Exposing the belly to another rat in a half-erect posture |
| Maintenance | Drinking | Drinking water |
| Maintenance | Eating | Eating food |
| Maintenance | Grooming | Self-grooming, when a rat is cleaning itself with rapid little nibbles |

#### 2.3.3. Faeces collection for corticosterone measurements

Faeces collection took place one day before and immediately after VBS housing. At the same time of the day, all rats were simultaneously housed in individual cages with food, water and clean bedding. They spent up to 4 h in their cages. Every 30 minutes, faeces produced were collected in microtubes and stored at −20°C until corticosterone extraction. Next, the samples were thawed and 0.1 g of faeces was added to 0.9 ml of 90% methanol, agitated for 30 min and centrifuged at 3000 rpm for 15 min. A 0.5 ml aliquot of the supernatant was added to 0.5 ml water; this extract was stored at −20°C. Corticosterone measurements were done with an enzyme immunoassay (EIA) following the method of Lepschy et al., [41] in the laboratory of Dr. Dehnhard at the Leibniz Institute of Zoo and Wildlife Research, Berlin. The antibody was purchased from Rupert Palme (University of Veterinary Medicine, Vienna, Austria), and has been described in detail in [42]. Briefly, a double antibody technique was used in association with a peroxidase conjugate, generating a signal quantitatively measurable by photometry. The concentration of corticosterone was expressed in µg per g of faecal material as an indicator of stress level in an individual. The change in corticosterone level (%) was calculated from the values obtained before and after the VBS.

#### 2.3.4. Social preference and recognition in the SRt

The protocol was adapted from Shahar-Gold et al., [43]. The test took place in a square open field (50 cm), with a small cage placed in one corner (Supplementary Fig. 6A). The intruder animals were older WH rats with a prior habituation to the procedure. A video camera placed above the open field was used to record the experiment. Each rat was tested on two consecutive days. On the first day, the subject was placed in the open field containing the empty small cage in a corner for a habituation period of 15 min (Hab). The intruder was then placed in the small cage, and the subject could freely explore the open field for 5 min (E1). Subsequently, the small cage with the intruder was removed from the open field, and the subject remained alone in the open field for 10 min. The encounter procedure was repeated two more times with the same intruder (E2, E3). On the second day, the first 15 min of habituation were followed by a fourth encounter (E4) of 5 min with the same intruder as on day 1. After this encounter, a break of 30 min took place, during which the subject remained alone in the open field. The last encounter then took place with an unfamiliar intruder placed in the same small cage for 5 min (Enew). The time spent in close interaction, including when the subject’s head was in contact with the grid or within 1 cm of the grid and the nose directed to the grid, was measured for each encounter (E1, E2, E3, E4 and Enew) and for the first 5 min of Hab. The social preference was calculated as the ratio of the interaction time in E1 and the interaction time during Hab. The short-term social recognition memory was calculated as the ratio of the interaction time in E1 and the interaction time in E3. The long-term social recognition memory was calculated by dividing the interaction time in Enew by the interaction time in E4.

#### 2.3.5. Exploration in the EPM

The apparatus (made of black painted wood) consisted of two open arms (50 cm x 15 cm), alternating at right angles with two closed arms enclosed by 40 cm high walls. The four arms opened onto a central area (15 cm x 15 cm). There was a small ridge along the edge of the open arms (1 cm wide). The whole maze was elevated 60 cm from the ground. A video camera mounted above the maze and connected to a computer outside the experimental room was used to observe and record animal’s behaviour. Light intensity in the open arms was 15 Lux.

The experimenter placed a rat in the central area of the maze facing a closed arm. The rat was allowed to freely explore the maze for 10 min. The time spent and entries in the open and closed arms were measured. Risk taking was evaluated as time and number of visits in the last third of the open arms, constituting the more risky areas [6].

### 2.4. Statistical analysis

R (3.5.1) and R studio (1.1.456) free softwares were used for the statistical analyses [44]. For each test, two levels of analysis were considered: first, the inter-strain comparison, where whole populations of WH *vs.* DA were compared, including INT animals; and second, the intra-strain comparison, where GDMs *vs.* PDMs were compared within each strain (excluding the INT animals).

Several non-parametric tests were used: a) the Fisher’s exact test was used to compare the number of GDM and PDM in WH and DA groups; b) the Wilcoxon sign test (RVAidememoire package) [45] was used to compare the performance of the animals to the indifference level (DDT, PDT and SRt); c) the Wilcoxon rank sum test was used to compare groups of animals (DA *vs.* WH, GDM *vs.* PDM, and cluster groups between them), and whenever appropriate a continuity correction was applied to the data with the Wilcoxon rank sum test; and d) the non-parametric ANOVA with permutation for repeated measures (lmPerm package) [46] was used to compare groups of animals along different time points. The one sample t-test was used to compare the performances with the indifference level in the RGT. For the global discrimination between strains, we used a random forest (RF) classification with leave-one out validation (randomForest package) [47]. The traits included in this analysis were the variables from the different tests. Seventeen traits were used: percentage of advantageous choices during the last 20 min (RGT score); flexibility score; mean latency to visit the feeder after a choice (latency RGT); AUC in DDT; activity in DDT; AUC in PDT; mean number of responses in FI; mean number of responses in EXT; activity in VBS housing; time open VBS; number of aggressive, affiliative and maintenance behaviours in VBS test; weight variation in VBS housing; corticosterone variation in VBS housing; social preference ratio; and short-term recognition ratio. Missing values (NA) were not tolerated by the model; therefore, some animals and variables had to be excluded from the analysis (for example, two animals did not produce faeces during faeces collection and the EPM was not included). n = 22 WH and n = 24 DA were included in the RF analysis.

## 3. Results

### 3.1. Cognitive and social abilities in DA and WH rats

#### 3.1.1. Decision-making abilities in the RGT

At the beginning of the test (first 10 min), rats of both strains chose the advantageous and disadvantageous options equally (Fig. 1B). After 10 min and until the end of the test, the average performance of the DA rats moved toward the most advantageous options (20 min: one sample t-test for DA: 0.95 CI [55, 76.6], p = 0.005), while the average performance of the WH rats remained at chance level for the entire duration of the test. However, at the end of the test (the last 20 min), large individual differences in choice became clear (Fig. 1C). In both strains, a majority of the rats preferred the most advantageous options at the end of the test (> 70% advantageous choices during the last 20 min of test; good decision makers or GDMs); a smaller proportion preferred the most disadvantageous options (< 30% advantageous choices; poor decision makers or PDMs) and a minority of the animals showed intermediate performance (INTs). Of the DA rats, 79% were GDMs (n = 31), 19% were PDMs (n = 8) and 5% were INTs (n = 2); of the WH rats, 50% were GDMs (n = 15), 40% were PDMs (n = 12) and 10% were INTs (n = 3). The proportion of GDMs, INTs and PDMs between strains were not statistically different (Fisher’s exact test, p=0.081), only the proportion of GDMs *vs.* non-GDMs (INTs and PDMs) was higher in the DA than the WH (Fisher’s exact test, p=0.04321). These observations could explain why the average performance of the DA rats was above the 50% indifference level while the WH rats were not. The development of choice preferences during the test of the GDMs on one hand and of PDMs on the other hand were similar between strains (Supplementary Fig. 1A).

In both strains, “fast” and “slow” GDMs could be identified (Supplementary Fig. 1B). In the DA rats, the majority of the GDMs were the “fast” type (76%; n = 23/30), choosing significantly and consistently the advantageous options at 20 min of testing. In the WH rats, only half of the GDMs were the “fast” type (53%, n = 8/15).

#### 3.1.2. Motivation for reward in the RGT

The latency to collect a reward after making a choice in the RGT was shorter in the WH rats (median 1.1 s) than in the DA rats (median 1.8 s; Fig. 1D, Wilcoxon rank sum test, W = 1151, p < 0.001). This difference was not due to the different proportions of GDMs and PDMs. In both strains, the PDM rats were faster than the GDM rats at collecting the reward (Fig. 1D, Wilcoxon rank sum test, WH: W = 147, p = 0.004; DA: W = 181, p = 0.047). Interestingly, the WH GDMs had the same latency as the DA PDMs (Fig. 1D).

#### 3.1.3. Cognitive flexibility in the reversed-RGT

The flexibility score indicates the propensity of an individual in the reversed-RGT to keep choosing (inflexibility) the same outcome as in the previous RGT or not choosing it (flexibility). All animals considered, DA and WH rats presented similar levels of cognitive flexibility (Fig. 1E; median 29% and 18% for DA and WH, respectively). In both strains and as expected for WH, all PDMs made highly inflexible choices in the reversed-RGT (low flexibility score; Fig. 1E). PDM rats kept choosing the hole(s) previously preferred (in the RGT), despite the outcomes of these choices now being different than in the RGT (Supplementary Fig. 2). In both strains, GDM rats had either high, intermediate or low flexibility scores (Fig. 1E). The proportion of GDMs with a high flexibility score (flexible GDMs) was 39% in DA and 33% in WH. Flexible GDMs progressively (trial after trial) switched their spatial preference from the nose-poke holes previously associated with the advantageous options (in the RGT) to the nose-poke holes currently associated with the advantageous options (Supplementary Fig. 2). 22% of DA GDMs and 20% of WH GDMs had no clear preference for either advantageous or disadvantageous options during the reversed-RGT. Finally, 39% of DA GDMs and 47% of WH GDMs showed an inflexible pattern of choices similar to the PDM rats (Fig. 1E) and kept choosing the hole(s) previously preferred in the RGT (Supplementary Fig. 2).

#### 3.1.4. Cognitive impulsivity in the DDT

In both strains, increasing the delay of delivering a highly palatable large reward decreased the preference for this option (Fig. 1F; Wilcoxon sign test, delay 0 s: DA 0.95 CI [85.1, 93.0], p < 0.001, WH 0.95 CI [81.4, 88.7], p < 0.001; delay 10 s: DA 0.95 CI [73.1, 93.0], p < 0.001). The sooner an individual rejects the large reward that is increasingly delayed, the more impulsive it is. On average, the DA rats preferred an immediate one-pellet reward over a delayed five-pellet reward when the delay reached 30 s (Wilcoxon sign test, 0.95 CI [19.4, 26.5], p < 0.001). Similarly, on average, WH rats preferred an immediate one-pellet reward over a delayed five-pellet reward when the delay reached 20 s (Wilcoxon sign test, 0.95 CI [10.8, 31.3], p = 0.001). Interestingly, although the preference for the high-reward option at a delay of 0 s was very strong in both strains (91% in DA and 84% in WH), the performance was significantly different between strains (Fig. 1F; Wilcoxon rank sum test with continuity correction, W = 891, p = 0.002). After normalizing performances to the preference at a delay of 0 s, the comparison of the AUC indicated that WH rats lost the preference for the high-reward option earlier than DA rats when the delay was added (Fig. 1G; Wilcoxon rank sum test, W = 923, p < 0.001). Within strains (and as expected for WH) [8], GDMs and PDMs had the same switching point and AUCs (Supplementary Fig. 3A and B).

#### 3.1.5. Cognitive risk taking in the PDT

In both strains, decreasing the probability of delivery of the most rewarding option (five pellets) also decreased the preference for this option (Fig. 1H; Wilcoxon sign test, probability 100%: DA 0.95 CI [73, 91.2], p < 0.001; WH 0.95 CI [80, 90], p < 0.001). A delivery probability of 20% for the five-pellet option is the point of indifference at which both options (certain – one pellet *vs.* uncertain – five pellets) are, on average, equivalent. If an animal prefers the certain option (one pellet) over the uncertain option (P = 80% to 20% – five pellets), it indicates an aversion to uncertainty. If an animal prefers the uncertain option (P = 20% to 9% – five pellets) over the certain option (one pellet), it indicates risk taking. DA rats lost their preference for the (uncertain) high-reward option when probability dropped to 17% (Wilcoxon sign test, 0.95 CI [50.8, 72.8], p = 0.063). WH rats lost their preference when probability dropped to 20% (Wilcoxon sign test, 0.95 CI [40.8, 66.7], p = 0.361). Comparison of the AUCs indicated that DA maintained a higher preference for the high reward with the decrease of reward probability than WH (Fig. 1I; Wilcoxon rank sum test, W = 516, p = 0.006). In both strains, the AUCs were comparable between GDMs and PDMs (Supplementary Fig. 3C and D).

#### 3.1.6. Anticipatory and perseverative behaviour in the FI-EXT schedule of reinforcement

DA anticipatory activity was higher, particularly during the first 20 s of the FI (Fig. 1J; non-parametric ANOVA with permutation, 1^st^ segment p < 0.001, 2^nd^ segment p = 0.004). The mean number of nose pokes was higher in DA rats than in WH rats for the 1 min FI (Fig. 1K; Wilcoxon rank sum test with continuity correction, W = 589.5, p = 0.039). DA rats nose poked more than WH rats during the 5 min EXT (Fig. 1M; Wilcoxon rank sum test with continuity correction, W = 690, p < 0.001), and this was the case during all the 1 min segments of EXT (Fig. 1L; non-parametric ANOVA with permutation, 1^st^ segment p = 0.002, 2^nd^ segment p = 0.045, 3^rd^ segment p < 0.001, 4^th^ segment p = 0.001, 5^th^ segment p = 0.015). Within strains, DA PDMs (n=7) nose poked significantly more than DA GDMs during EXT (Supplementary Fig. 4B; Wilcoxon rank sum test with continuity correction, W = 35, p = 0.043); however, this was not observed in WH.

#### 3.1.7. Natural behaviours expressed in the VBS

In both strains, the behaviours most frequently observed in the VBS were huddle, eating and struggle at feeder (with median number of occurrences > 5 in 100 30 s videos on the last two days of VBS housing; Fig. 2A). The 19 other scored behaviours (allogrooming, attending, drinking, grooming, aggressive grooming, attack, embracing, fight, following, mounting, mutual upright posture, pinning, struggle at water, struggle in tunnel, flight, freezing, lateral defence, supine posture and upright defence) were seen more rarely (median number of occurrences < 5 in 100 30 s videos on the last two days of VBS housing) and are grouped in the composite category “19 others” in Figure 5 (for further details, see Supplementary Fig. 5A). Considering the three most frequent behaviours, DA rats huddled more and struggled at the feeder less than WH rats (Fig. 2B; Wilcoxon rank sum tests with continuity correction, huddle: W = 984, p < 0.001; struggle at feeder: W = 313.5, p = 0.005). Strains did not differ in their number of bouts of eating. The occurrences of huddle, eating and struggle at feeder were similar between PDMs and GDMs in both strains (Supplementary Fig. 5B).

**Figure 2.**
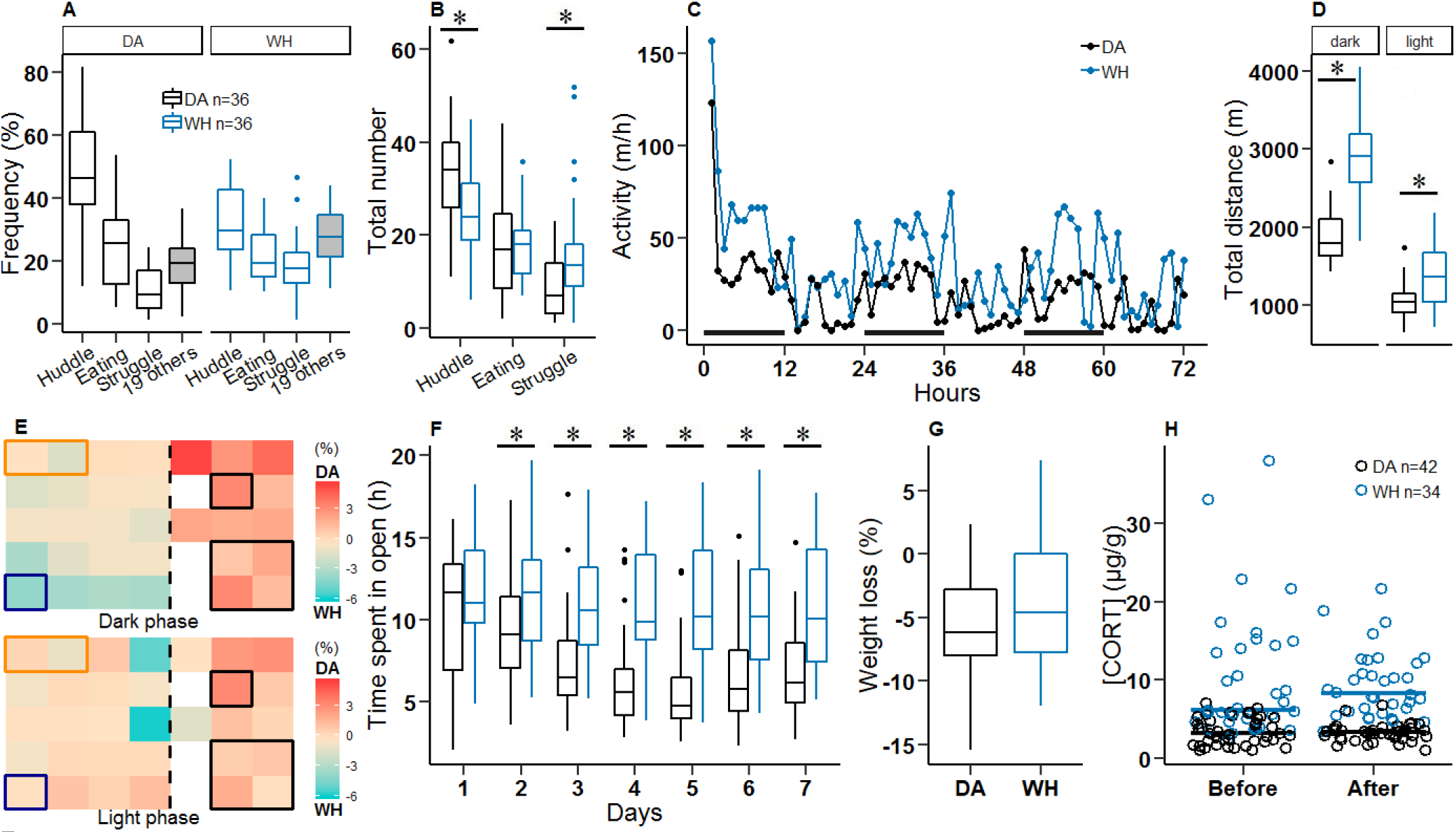
Daily activity, behavioural and biological measures of Dark Agouti (DA) and Wistar Han (WH) rats during the Visible Burrow System (VBS) housing. **A** Relative frequency of occurrence of behaviours in the VBS. White boxes represent a unique type and grey boxes represent a composite behaviour category. “Struggle” = “struggle at feeder”. “19 Others” comprised the 19 behaviours (all behaviours minus the three main behaviours) scored during the VBS video analysis but which had a median < 5 in each strain due to their rare occurrence. **B** Occurrence of the three main types of behaviours observed in the VBS (50 min observation). **C** Typical locomotor activity of one DA and one WH individual during the first three days in the VBS. Bars indicate dark phase. **D** Total distance travelled during the dark and light phases over seven days in the VBS. **E** Difference in place preference (%) between DA and WH during the dark and light phases over seven days of VBS housing. Red indicates a preference of the DA relative to WH for each of the 32 zones of the VBS (corresponding to the 32 RFID detectors located beneath the VBS cage). Rectangles indicate the locations of feeder (orange), water bottle (blue), and small and large chambers in the burrow area (black). The vertical dashed line indicates the separation between the open area (left side) and the burrow area (right side). **F** Total time spent in the open area. **G** Weight loss after VBS housing. **H** Concentration of corticosterone in faeces before and after VBS housing. Horizontal bar: median of each group. DA in black and WH in blue; Panels A-G: WH, n = 36; DA, n = 36 and panel H: WH, n = 34; DA, n = 42. * p < 0.05, DA *vs.* WH, Wilcoxon rank sum test except panel F ANOVA with permutations for repeated measures. The VBS test was conducted with n = 6 individuals in the cage at a time.

#### 3.1.8. Total distance travelled in the VBS

Both DA and WH rats changed their activity (i.e., the distance travelled) with the light/dark phase (Fig. 2C). Both strains were more active during dark phases (Fig. 2C). Over all days, locomotion in WH rats was higher than in DA rats during both dark and light phases (Fig. 2D; dark phase: Wilcoxon rank sum test, W = 45, p < 0.001; light phase: Wilcoxon rank sum test, W = 313, p < 0.001). During the dark phase, the WH PDMs were more active than the WH GDMs (Supplementary Fig. 5C; Wilcoxon rank sum test, W = 60, p = 0.005).

#### 3.1.9. Place preference in the VBS

DA rats preferred to stay in the burrow area significantly more than WH rats, both during the dark phase (Fig. 2E, top panel; Wilcoxon rank sum test, W = 105, p <0.001) and during the light phase (Fig. 2E, bottom panel; Wilcoxon rank sum test, W = 371, p = 0.001). Furthermore, during the light phase, WH rats were mostly present in the entry zones of the burrow area (Fig. 5E). The WH GDMs preferred staying in the burrow more than the WH PDMs during the dark phase (Supplementary Fig. 5D; Wilcoxon rank sum test, W = 195, p = 0.038) and the same tendency was observed in DA rats (Supplementary Fig. 5D).

#### 3.1.10. Total time spent in the open area of the VBS across days

The DA rats spent less time in the open area starting from day 2 (non-parametric ANOVA with permutation, day 2 p = 0.030) than WH rats (Fig. 2F). There was no difference in the time spent in the open area across day between DA GDMs and DA PDMs, whereas in WH the PDMs tended to spend more time in the open than GDMs starting on day 3 (Supplementary Fig. 5E).

#### 3.1.11. Weight loss during VBS housing

Before being housed in the VBS (and in general), DA rats were smaller and lighter than WH rats (Supplementary Fig. 5F; Wilcoxon rank sum test with continuity correction W = 0, p < 0.001). During their stay in the VBS, DA and WH rats lost the same relative weight (Fig. 2G). However, DA GDMs lost more weight than DA PDMs (Supplementary Fig. 5G; Wilcoxon rank sum test with continuity correction, W = 35, p = 0.039).

#### 3.1.12. Corticosterone (metabolite) levels after VBS housing

At baseline (before the VBS housing), the concentration of corticosterone in DA rats was lower than in WH rats (Fig. 2H; Wilcoxon rank sum test W = 206, p < 0.001). After VBS housing, the corticosterone levels in DA and WH rats were unchanged (Fig. 2H). In both strains, corticosterone levels were not different between GDMs and PDMs, either before or after VBS housing (Supplementary Fig. 5H).

#### 3.1.13. Social preference and social recognition memory in the SRt

In the SRt, both strains exhibited a clear preference for social *vs.* non-social cues and an accurate short-term social recognition memory. Rats spent more time exploring the unfamiliar social partner during the encounter 1 (E1) than an unfamiliar non-social cue (empty box) during the habituation phase (Hab, Fig. 3A; Wilcoxon rank sum test with continuity correction, WH: W = 576, p < 0.001; DA: W = 258.5, p < 0.001). Exploration time was twice as long in E1 as in Hab (Fig. 3B; social preference ratio E1/Hab >1, Wilcoxon sign test DA: 0.95 CI [1.3, 2.8], p = 0.030 and WH: 0.95 CI [2.2, 3.4], p < 0.001). WH rats had a higher social preference ratio than DA rats (Wilcoxon rank sum test, W = 121, p = 0.016). The third time WH and DA rats encountered the same animal (E3), the time spent exploring this animal was significantly reduced compared to their first encounter (E1), indicating effective short-term social recognition memory (Fig. 3A; Wilcoxon rank sum test with continuity correction, WH: W = 484.5, p < 0.001; DA: W = 225, p = 0.018). Due to experimental limitations, long-term social recognition memory could not be evaluated, although it is likely that both strains did have such memory (Supplementary Fig. 6A). In both strains, the social preference ratio and short-term memory ratio did not differ between GDMs and PDMs (Supplementary Fig. 6C and D).

**Figure 3.**
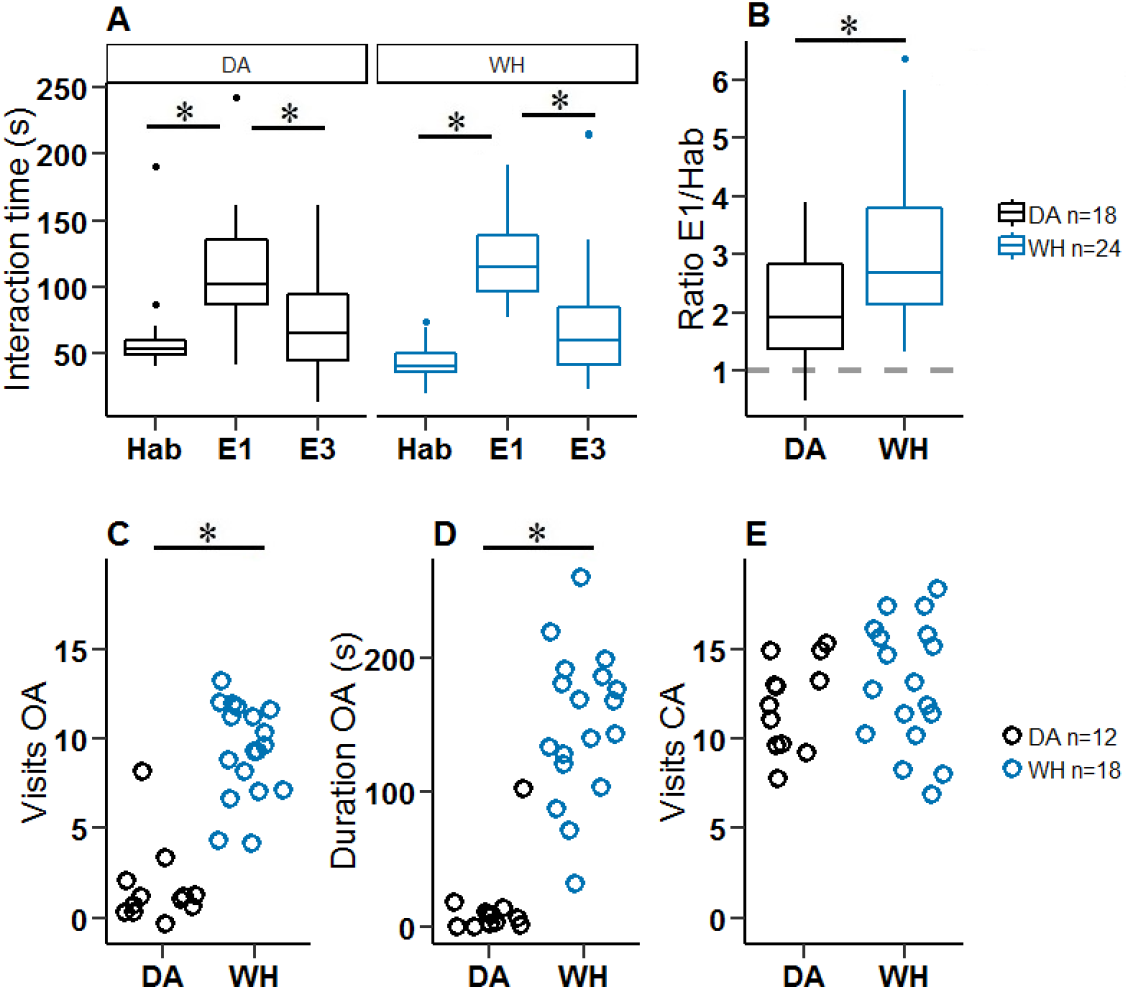
Social preference, social short-term recognition and exploration of the EPM in Dark Agouti (DA) and Wistar Han (WH) rats. **A** Interaction times during the social recognition test. Hab: non-social cue (empty box) present during the habituation phase; E1: first encounter with intruder (unfamiliar); E3: third encounter with same intruder (familiar); Wilcoxon rank sum test Hab *vs*. E1 and E1 *vs.* E3. **B** Social preference represented as the ratio of exploration times in E1 and in Hab, DA *vs.* WH (Wilcoxon rank sum test). **C** Total number of visits to the open arms (OA), DA in black and WH in blue. **D** Time spent in the OA, DA *vs.* WH (Wilcoxon rank sum test). **E** Total number of visits to the closed arms (CA). Maximum exploration time was 10 min. DA in black and WH in blue, * p < 0.05.

#### 3.1.14. Exploration in the EPM

DA rats expressed very different behaviour in the EPM compared to WH rats. DA rats very rarely (or never) visited the open arms of the maze (Fig. 3C; Wilcoxon rank sum test with continuity correction, W = 5.5, p < 0.001) and for a very short time (Fig. 3D; Wilcoxon rank sum test with continuity correction, W = 3, p < 0.001) compared to WH rats. Only one DA individual visited the part of the maze that was furthest from enclosing walls (the last third of the open arms), as opposed to all the individuals in WH (data not shown). DA and WH rats had the same number of visits to closed arms (Fig. 3E). Within strains, no differences were observed between PDMs and GDMs for the parameters of total number of visits to open arms, total time spent in open arms or total number of visits to the last third of the open arms (Supplementary Fig. 7).

#### 3.1.15. Inter-individual differences within DA and WH

In both strains, GDMs and PDMs showed similar tendencies in all tests (see Table 2 for details). In both strains, PDMs were faster to collect the reward than GDMs in the RGT, and all showed higher cognitive inflexibility in the reversed-RGT. In the VBS, the WH PDMs were more active during the dark phase, did not prefer the burrow area during the dark phase and spent more time in the open area on day 4 than the WH GDMs. In the VBS, the DA PDMs lost less weight than the DA GDMs (Table 2).

**Table 2:**
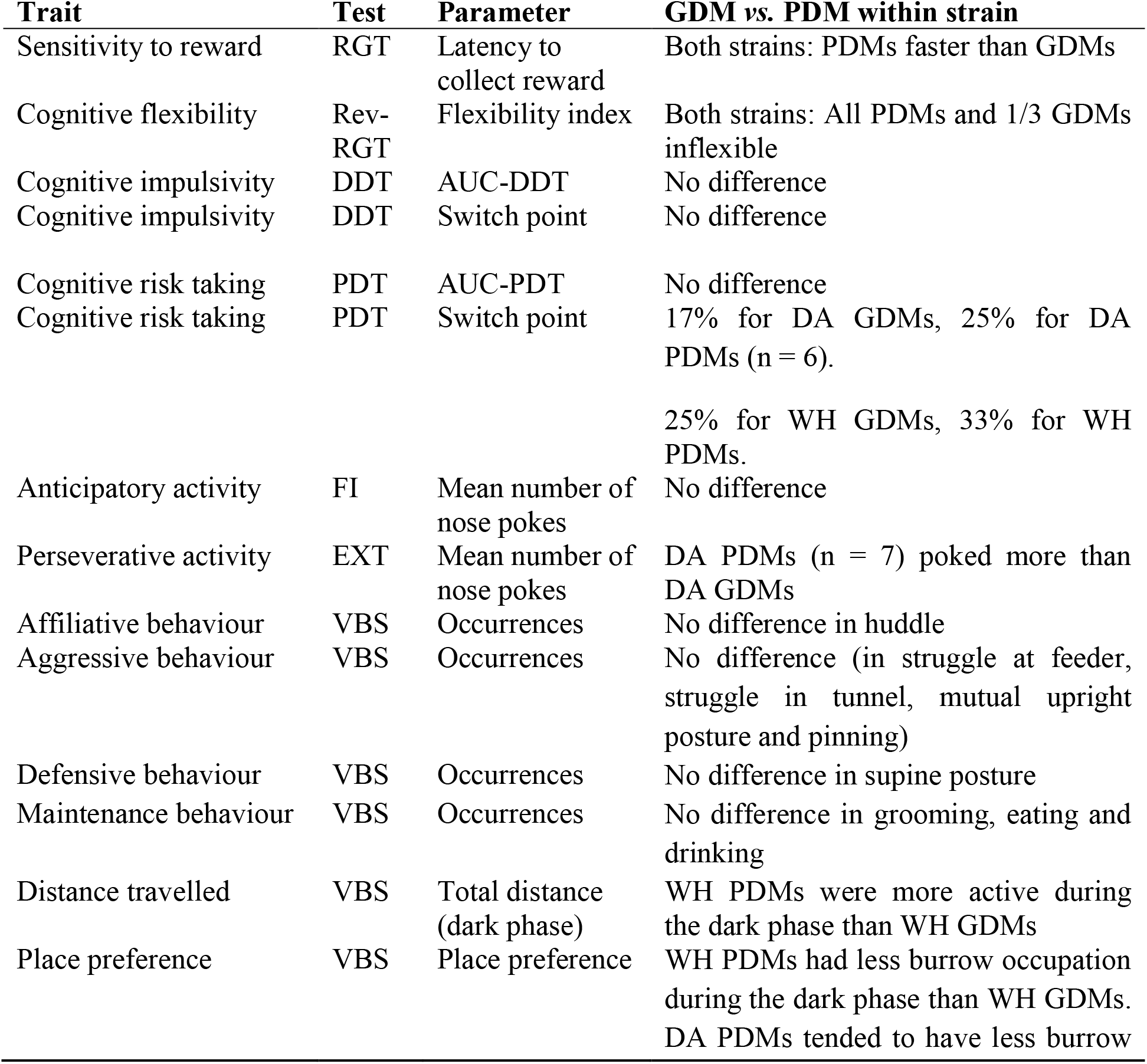
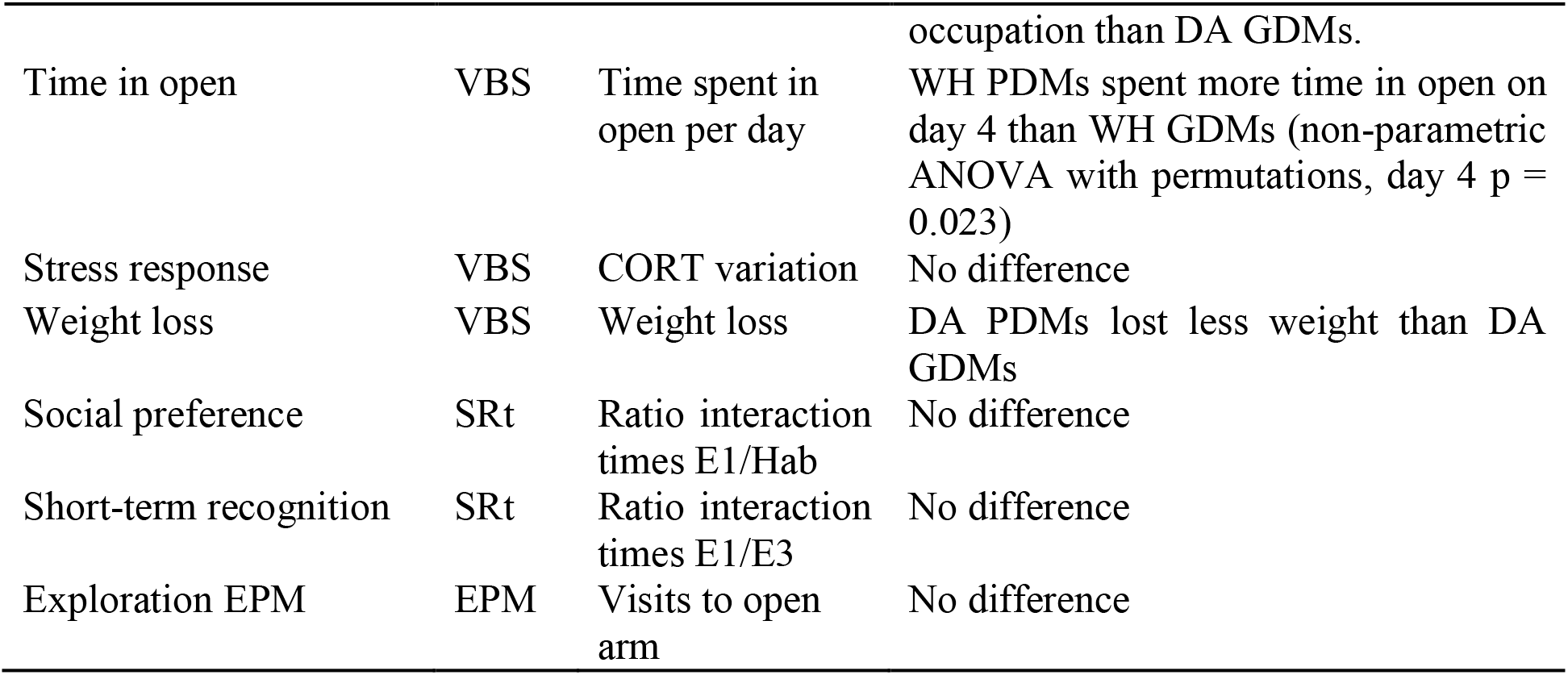
Behaviours of the GDMs and PDMs in DA and WH strains.

### 3.2. Identification of the key variables discriminating WH from DA strain

We performed an RF classification with a leave-one-out cross-validation (LOOCV) to quantify the efficiency of each of the previously described cognitive and social functions to distinguish WH and DA strains from each other. The RF was run using the behavioural and biological variables described above (Refer to the Methods section for a description of adjustment of measures and variables due to missing values). In brief, the decision trees of the RF with LOOCV led to the prediction of the strain of each of a given individual by comparing its performance (for each variable) to the performance of the other individuals for which the strain was known. For WH and DA variables, the prediction of the strain was high, with an accuracy of 84% (±0.72 SD over 10 runs). The importance of each variable to accurately differentiate the strains was given by the Gini index of the RF (Fig. 4A). The most discriminating variables were the AUC of the DDT and the distance travelled in the VBS (Gini index > 3), followed by the latency to collect a reward in the RGT and the total time spent in the open area in the VBS (3 > Gini index > 2; Fig. 4A). Of lesser significance were the social preference index in the social preference test, the AUC of the PDT, and the number of aggressive behaviours in the VBS (2 > Gini index > 1). The least discriminating variables were the total number of affiliative behaviours, the weight variation and the total number of maintenance behaviours in the VBS; the mean number of responses during FI and EXT; the decision-making score in the RGT; the variation of CORT levels; the short-term recognition memory in the social recognition test; the total activity in the DDT; and the flexibility score in the reversed-RGT (Gini index < 1; Fig. 4A). As an example, an RF classification including the two most discriminating variables (the distance travelled in the VBS and the AUC of the DDT) attributed the correct strain to 41 rats out of a total of 46 rats (Fig. 4B). On the contrary, an RF including only the variables with a Gini index < 1 resulted in a drop in accuracy to 50% (chance level, not shown).

**Figure 4.**
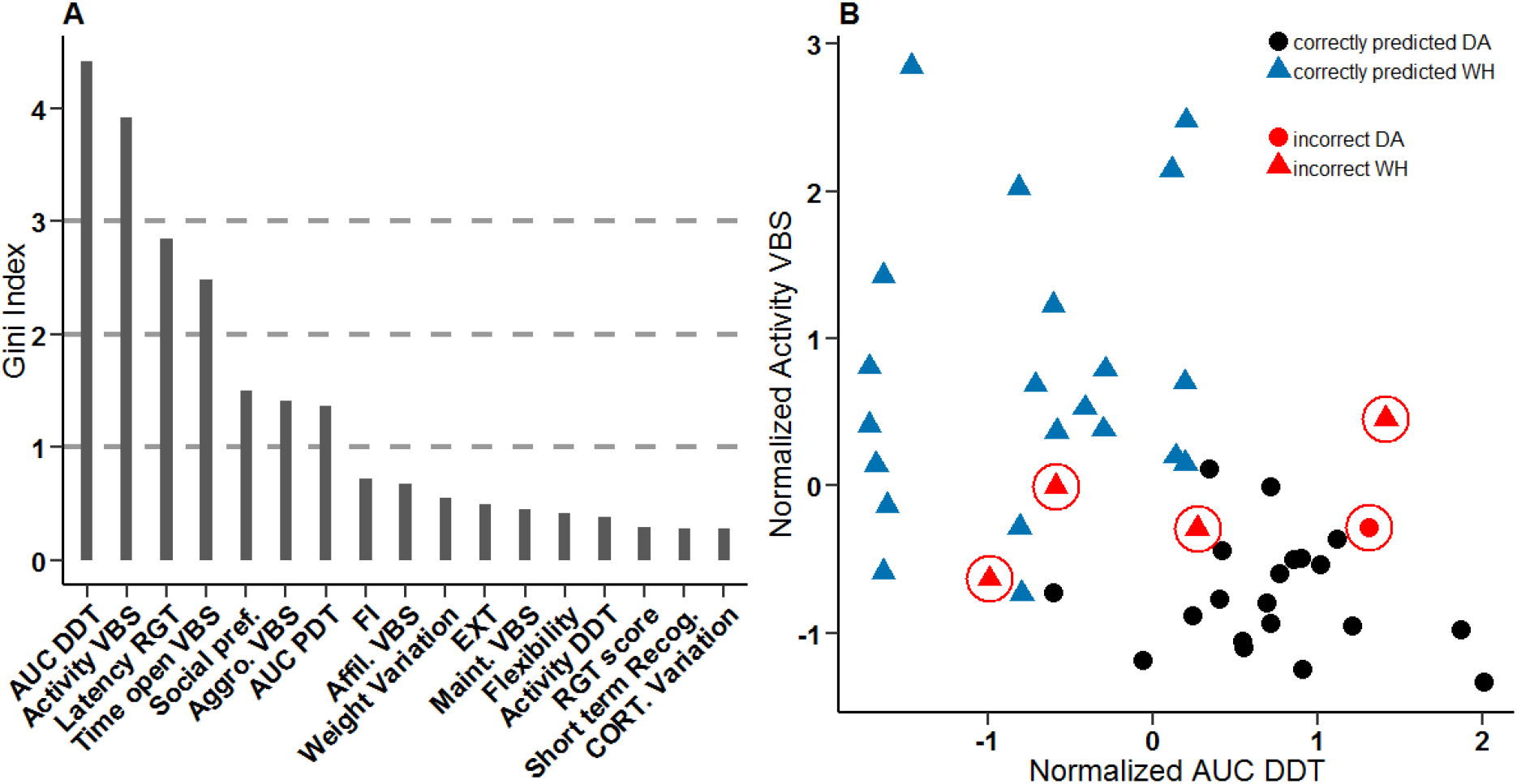
Discriminating classification of the DA and WH. **A** Gini index for each trait used for the random forest (RF) classification. Dashed lines are included to sort the variables in groups of importance. pref. = preference, Aggro. = aggressive, affil. = affiliative, maint. = maintenance, CORT = corticosterone. **B** RF classification for the two most discriminating variables. DA, n = 24, in black; WH, n = 22, in blue. Symbols show predicted strain by the RF. DA: dot, WH: triangle. Red circles indicate an incorrect prediction.

## 4. Discussion

### 4.1. Behavioural performance of PDMs and GDMs from DA and WH strains

One of the advantages of the RGT is the possibility it offers to uncover which decision-making strategy each individual of a healthy population of rats will spontaneously use to cope with complex and uncertain choice options. Here we found that, similar to WH, each individual DA could be classified in one of the three typical categories. We identified GDM strategists, which secured more food over the long term, although they earned smaller amount of food in each trial; PDM strategists, which secured less food over the long term, although they earned larger amounts of food in each trial but were penalized by long waiting periods; and INT individuals, which seemed indifferent to reward options. Although not significant, the higher number of GDMs found in the DA rats compared to the WH rats could explain their more advantageous performance as a group (averaged performance) during the RGT compared to the WH, which on average stayed at chance level for the entire duration of the test. In a follow-up study, we will evaluate the effect of a lack of central 5-HT on the animals’ decision-making abilities in the RGT. Thus, the large number of GDMs in healthy individuals will help us to quantify the effect of this genetic manipulation, which is expected to shift the behavioural profile from GDM to PDM.

Interestingly and independent of strain, we found that all GDM and PDM rats behaved as expected with regard to their decision-making type in the reversed-RGT and in anticipation of rewards (test of reward sensitivity in the RGT) [6]. All PDMs of either strain rapidly and steadily chose the least advantageous options in the long term in the RGT; they were more sensitive to the reward than GDMs and were unable to flexibly adjust their behaviour during the reversed-RGT. For humans, a new computational modelling of the analogous Iowa Gambling Task called Outcome-Representation Learning predicts that poor decision making of drug users could be due to higher reward sensitivity and more exploratory behaviour (in cannabis users), lower punishment sensitivity (in abstinent heroin users) and higher inflexibility perseverance (in abstinent amphetamine users) [48]. The expression of the same key features between PDMs of genetically distinct strains of rats and, to a certain extent, to results found in humans [49,50] suggests a strong conservation of this potential endophenotype within and between species.

As seen in previous studies in WH rats but now also in DA rats, the GDMs were not a single homogeneous group of rats [5,8]. While some (50% to 75%) were faster than others in choosing the advantageous options during the RGT (at only 20 min of test), in the reversed-RGT only one-third of GDMs were able to flexibly adjust their behaviour.

In addition, differences were not observed between PDMs and GDMs in either strain in cognitive impulsivity (DDT) or risk-based decision-making tests (PDT). Although the result of the DDT was expected [8], the lack of difference in the PDT between PDMs and GDMs was more surprising. Indeed, in another version of the RGT (i.e., the rGT, with a testing phase lasting three days and two options only (a reward, given as sweet pellets, or a punishment, given as quinine pellets)), poorer decision-making abilities were correlated with higher preferences in the PDT for the risky (large reward, uncertain outcome) options [24]. The differences between the experimental procedures of each study (the protocols of the RGT/rGT and the PDT were equivalent, but not identical) and in the definition of what constituted poor decision making (in RGT, spontaneous healthy PDMs were different from GDMs; in rGT, all rats were “GDMs”, but some individuals made poorer decisions than others) may be the reasons for the discrepancies between these results. However, it is noteworthy that in the human literature a loss of control over risk (probability)-based choices is not characteristic of all PDM-associated psychiatric disorders. Patients with pathological gambling [51], alcohol dependence [52], schizophrenia [53] and autism [54] are more risky decision makers than patients with obsessive-compulsive disorder [55], pathological buying disorder, Huntington’s disease [56] or suicidal attempts [57]. These and our results indicate that preference for high-risk (probabilistic) options may be a marker of pathology rather than a marker of vulnerability to diseases and thus may be preferentially observed in “ill-induced” PDMs than in healthy PDM rats.

In the FI-EXT test, we only witnessed increased motor impulsivity in DA PDMs during EXT, and did not witness this in either FI or EXT in WH. This inconsistent result in WH rats compared to our previous study may be due to the use of a different manipulandum (nose-poke holes instead of levers) for the operant response [8]. It is also possible that for WH rats, repetitive nose poking in a hole was too physically demanding to exhibit anticipatory or perseverative behaviours compared to pressing a lever. Very few studies have investigated the consequences of this difference in operant responding. Although Mekarski [58] defended nose poking to be a more innate behaviour than lever pressing, it has also been shown that escalation behaviour is better achieved with lever pressing and not nose poking in mice [59]. We also explored if PDM and GDM rats differed in their social skills. In the VBS, compared to GDM rats, PDM rats expressed a higher level of activity, less occupation of the burrow during dark phases, longer time spent in the open area of the cage (WH PDMs), and limited weight loss (DA PDMs). In the VBS, these features characterize dominance in rats (along with the number and location of wounds, which were not witnessed in this study) [60], suggesting a more dominant status for PDM rats than for GDM rats. In the same line, Davis et al., [61] found that individual dominance correlated with higher motivation for rewards and higher exploration of risky zones in the EPM. These are also two known characteristics of PDM rats [6]. Interestingly, PDMs were not more aggressive or less affiliative in the VBS than GDMs and presented a similar interest for the social cue in the SRt. While the experimental measurement of dominance in rats is often reduced to a one-time measure of aggression level (i.e., the resident intruder paradigm), Buwalda et al. [62] showed that the level of aggression in the resident-intruder paradigm and in the VBS were not correlated with dominance. However, a more realistic view of dominance should consider its multidimensional features including privileged access to resources [63,64], lower sensitivity to stressors [65] and non-agonistic behaviours [66]. Indeed, social hierarchy is a dynamic feature that depends on the outcome of each type of interaction [66,67]. In humans, excessive aggression is a disruptive symptom widely distributed among psychiatric disorders. Studies have shown that decision making and aggression-related behaviours could share biological markers, such as MAO A, SERT, TPH1 and TPH2 proteins [68,69]. In further studies, we will use the rich semi-natural and around-the-clock experimental conditions of our VBS housing to explore more specifically which social domains and how social hierarchy develop along with decision-making abilities and serotonin manipulations.

The reproducibility and conservation of the socio-cognitive and behavioural phenotypes of GDM and PDM individuals in the two genetically different strains of WH and DA rats support a good translational validity of these complex phenotypes, not only between strains but likely also between species (e.g., rats and humans). Following the Research Domain Criteria framework (RDoC), which promotes the exploration of cross-species endophenotypes for better translational value of preclinical studies [70,71], this study presents the PDM rats as a promising animal model for the identification of the specific biological circuits underlying equivalent patterns of deficits which could be observed in patients (or healthy relatives) and independently of their disorders’ categories. Both DA and WH rat strains offer interesting individual variations in behaviour, allowing the use of both strains for the study of the underlying mechanisms of poor decision making and associated disorders. It will be possible to examine the risk factors responsible for the transition from vulnerability to pathology by comparing the expression of each of the PDM-associated traits and how the neural substrates of this phenotype overlap or differ in ill-induced *vs.* healthy PDMs.

### 4.2. Strain differences between DA and WH

Beside the inter-individual differences within strains, we found at the group level that WH rats were, on average, more sensitive to reinforcement and more impulsive in the DDT, but less prone to take risks in the PDT compared to DA rats. In the DDT and PDT, WH rats dismissed both the delayed and uncertain option more rapidly than the DA rats in favour of the immediate or certain option, although this meant that the option associated with the largest reward (absolute value) was abandoned for a one-pellet option. The discounting factor (delay or probability) appeared to have a stronger impact on the subjective evaluation of rewards by WH rats, and WH rats had a lower tolerance to uncertain situations when rewards were involved compared to DA rats. In the VBS, WH rats were more aggressive, more active (higher distance travelled) and spent more time in the open area of the VBS than DA rats.

In biomedical research, the WH line is one of the two most commonly used strains of rats (the other being Sprague Dawley) [72]. This research included studies investigating reward-related disorders such as drug addiction [73,74] and poor impulse control-related disorders such as substance abuse, eating disorders, ADHD or manic disorders [75,76]. WH rats are also used in studies on reward processing and valuation [77], and have been found to have a high tendency for compulsive and impulsive behaviours [78,79].

In contrast, DA rats made more perseverative responses in the FI-EXT test in anticipation of a reward and during extinction phases, indicating either a lower tolerance to frustrating inactive phases of the test or higher motor impulsivity compared to WH. Knowing that the conditions for this test may not have been optimal (as the low level of activity may be due to the requirement for nose-poke holes instead of lever presses) and that such higher motoric response was not similarly observed in the training phase of the DDT (as both variables are correlated) [8], we prefer not to place too much emphasis on this result. Finally, DA rats were more affiliative in the VBS, preferred hiding in the burrows and were more fearful of the open arms of an EPM. They also had a weaker social preference in the SRt, which could be due to the avoidance of the centre of the open field during the first 5 min of habituation in this test. These results could confirm a specific fear of the elevated and widely open spaces, as discussed elsewhere [80,81].

With DA rats presenting a more compulsive, anxious and prosocial phenotype, this strain seems promising for studies on anxiety-related disorders. For example, patients diagnosed with anxiety disorder are extremely fearful/anxious of real-life threats (as opposed to unreal life-threatening concerns of OCD patients); they can express un-ritualized compulsive behaviours and, in the case of social anxiety disorder (social phobia), a subcategory of anxiety disorder, they show strong social contact avoidance and/or seek to reduce their social fear (DSM-5) [82]. Anxiety indeed appear to be a trait often witnessed in inbred lines of mice [33]. Finally, and despite their remarkable differences, DA and WH rats also shared similar traits. For example, they presented higher levels of huddling, eating and struggling at the feeder than other behaviours during VBS housing, and equivalent corticosterone level and weight loss after VBS housing.

### 4.3. Prediction of the strain differences with RF analysis

Although we identified specific traits on which DA and WH strains spontaneously differed in performance, using a RF classification method helped to determine which of these traits were more characteristic of one strain than the other. These were the ability to wait for a reward in the DDT, the motivation to collect a reward in the RGT, and the level of activity and time spent in the open area of the VBS. The RF classifier was less able to accurately differentiate strains based on the expression of their affiliative and maintenance behaviours, weight variation, decision making or flexibility. The RF classification results were similar to those obtained after a principal component analysis (Supplementary Fig. 8A and D).

In other words, the most critical difference between WH and DA rats related to behavioural control when facing a (delayed or non-delayed) reward as seen in the DDT (cognitive impulsivity) and the RGT (reward seeking), respectively. Based on this observation, it could also be argued that the increased time the WH rats spent in the open area of the VBS was driven by the presence of the only food source of the cage being in this area, although this zone was also potentially the most aversive zone of the cage.

## 5. Conclusion

In this study, we compared several abilities of DA and WH rats at the group and the individual levels using multiple cognitive tests, a social naturalistic set-up and assays of physiological responses.

Both the dimensional and group approaches provided new insights for the preferential use of each strain in future neuropsychopharmacological studies and further advanced our knowledge of the complex phenotype of the healthy PDM and GDM. At the group level, we identified specific traits on which these genetically distinct strains spontaneously differed the most (AUC of the DDT, distance travelled in the VBS, latency to collect a reward in the RGT and total time spent in the open area in the VBS). The WH and DA strains could preferentially be used to model reward sensitivity and impulsivity on one side and compulsivity and anxiety-related behaviours on the other side.

At the individual level, we could reproduce previous findings in WH rats and generalize them to the DA strain. Each PDM individual of either strain displayed a similar naturally occurring combination of behavioural traits, including a higher sensitivity to reward, higher cognitive inflexibility and higher social rank, but no cognitive impulsivity in delay- or probability-based decision-making tasks, no deficits in social recognition and no differences in corticosterone response to stressors. The multidomain profile of the PDM individuals should be suitable to reveal bio-behavioural specificities highly relevant for the study of human mental illnesses. In a follow-up study, we will directly interfere with rats’ central serotonergic system and evaluate the impact of this intervention in the concomitant modulation of the PDM-associated traits.

## Supporting information

Supplementary results

## Acknowledgements

We want to thank Patrik Bey, Melissa Long, Alexej Schatz, Dr. Martin Dehnhard and his team and the FEM team for their technical assistance and our colleagues of the Winter lab who made insightful comments on a previous version of the manuscript.

## References

[1] F. Dellu-Hagedorn, M. Rivalan, A. Fitoussi, P. De Deurwaerdère, Inter-individual differences in the impulsive/compulsive dimension: deciphering related dopaminergic and serotonergic metabolisms at rest, Philos. Trans. R. Soc. Lond. B. Biol. Sci. 373 (2018). doi:10.1098/rstb.2017.0154.

[2] E.J. Nestler, S.E. Hyman, Animal models of neuropsychiatric disorders, Nat. Neurosci. 13 (2010) 1161–1169. doi:10.1038/nn.2647.

[3] M. Rivalan, C. Blondeau, F. Dellu-Hagedorn, Chapter2. Modeling symptoms of mental disorders using a dimensional approach in the rat., Loop. (2009). http://loop.frontiersin.org/publications/41109606.

[4] B. Cao, J. Wang, M. Shahed, B. Jelfs, R.H.M. Chan, Y. Li, Vagus Nerve Stimulation Alters Phase Synchrony of the Anterior Cingulate Cortex and Facilitates Decision Making in Rats, Sci. Rep. 6 (2016) 35135. doi:10.1038/srep35135.

[5] A. Fitoussi, C. Le Moine, P. De Deurwaerdère, M. Laqui, M. Rivalan, M. Cador, F. Dellu-Hagedorn, Prefronto-subcortical imbalance characterizes poor decision-making: neurochemical and neural functional evidences in rats, Brain Struct. Funct. 220 (2015) 3485–3496. doi:10.1007/s00429-014-0868-8.

[6] M. Rivalan, S.H. Ahmed, F. Dellu-Hagedorn, Risk-prone individuals prefer the wrong options on a rat version of the Iowa Gambling Task, Biol. Psychiatry. 66 (2009) 743–749. doi:10.1016/j.biopsych.2009.04.008.

[7] M. Rivalan, E. Coutureau, A. Fitoussi, F. Dellu-Hagedorn, Inter-Individual Decision-Making Differences in the Effects of Cingulate, Orbitofrontal, and Prelimbic Cortex Lesions in a Rat Gambling Task, Front. Behav. Neurosci. 5 (2011). doi:10.3389/fnbeh.2011.00022.

[8] M. Rivalan, V. Valton, P. Seriès, A.R. Marchand, F. Dellu-Hagedorn, Elucidating Poor Decision-Making in a Rat Gambling Task, PLoS ONE. 8 (2013) e82052. doi:10.1371/journal.pone.0082052.

[9] R. van den Bos, W. Davies, F. Dellu-Hagedorn, A.E. Goudriaan, S. Granon, J. Homberg, M. Rivalan, J. Swendsen, W. Adriani, Cross-species approaches to pathological gambling: a review targeting sex differences, adolescent vulnerability and ecological validity of research tools, Neurosci. Biobehav. Rev. 37 (2013) 2454–2471. doi:10.1016/j.neubiorev.2013.07.005.

[10] B.N. Cuthbert, Research Domain Criteria: toward future psychiatric nosologies, Dialogues Clin. Neurosci. 17 (2015) 89–97.

[11] A.V. Kalueff, A.M. Stewart, C. Song, I.I. Gottesman, Targeting dynamic interplay among disordered domains or endophenotypes to understand complex neuropsychiatric disorders: Translational lessons from preclinical models, Neurosci. Biobehav. Rev. 53 (2015) 25–36. doi:10.1016/j.neubiorev.2015.03.007.

[12] H.G. Baumgarten, Z. Grozdanovic, Psychopharmacology of central serotonergic systems, Pharmacopsychiatry. 28 Suppl 2 (1995) 73–79. doi:10.1055/s-2007-979623.

[13] S. Enge, M. Fleischhauer, K.-P. Lesch, A. Reif, A. Strobel, Serotonergic modulation in executive functioning: linking genetic variations to working memory performance, Neuropsychologia. 49 (2011) 3776–3785. doi:10.1016/j.neuropsychologia.2011.09.038.

[14] D. Kiser, B. Steemers, I. Branchi, J.R. Homberg, The reciprocal interaction between serotonin and social behaviour, Neurosci. Biobehav. Rev. 36 (2012) 786–798. doi:10.1016/j.neubiorev.2011.12.009.

[15] D. Mendelsohn, W.J. Riedel, A. Sambeth, Effects of acute tryptophan depletion on memory, attention and executive functions: A systematic review, Neurosci. Biobehav. Rev. 33 (2009) 926–952. doi:10.1016/j.neubiorev.2009.03.006.

[16] T.W. Robbins, A.F.T. Arnsten, The neuropsychopharmacology of fronto-executive function: monoaminergic modulation, Annu. Rev. Neurosci. 32 (2009) 267–287. doi:10.1146/annurev.neuro.051508.135535.

[17] J. Waider, N. Araragi, L. Gutknecht, K.-P. Lesch, Tryptophan hydroxylase-2 (TPH2) in disorders of cognitive control and emotion regulation: a perspective, Psychoneuroendocrinology. 36 (2011) 393–405. doi:10.1016/j.psyneuen.2010.12.012.

[18] C.A. Winstanley, The utility of rat models of impulsivity in developing pharmacotherapies for impulse control disorders, Br. J. Pharmacol. 164 (2011) 1301–1321. doi:10.1111/j.1476-5381.2011.01323.x.

[19] S.N. Young, M. Leyton, The role of serotonin in human mood and social interaction. Insight from altered tryptophan levels, Pharmacol. Biochem. Behav. 71 (2002) 857–865.

[20] A.R. Maher, G. Theodore, Summary of the comparative effectiveness review on off-label use of atypical antipsychotics, J. Manag. Care Pharm. JMCP. 18 (2012) S1–20.

[21] C.B. Nemeroff, Psychopharmacology of affective disorders in the 21st century, Biol. Psychiatry. 44 (1998) 517–525.

[22] E. Hollander, J. Rosen, Impulsivity, J. Psychopharmacol. Oxf. Engl. 14 (2000) S39–44. doi:10.1177/02698811000142S106.

[23] J.R. Homberg, R. van den Bos, E. den Heijer, R. Suer, E. Cuppen, Serotonin transporter dosage modulates long-term decision-making in rat and human, Neuropharmacology. 55 (2008) 80–84. doi:10.1016/j.neuropharm.2008.04.016.

[24] S. Koot, F. Zoratto, T. Cassano, R. Colangeli, G. Laviola, R. van den Bos, W. Adriani, Compromised decision-making and increased gambling proneness following dietary serotonin depletion in rats, Neuropharmacology. 62 (2012) 1640–1650. doi:10.1016/j.neuropharm.2011.11.002.

[25] C.A. Winstanley, J.W. Dalley, D.E.H. Theobald, T.W. Robbins, Fractionating impulsivity: contrasting effects of central 5-HT depletion on different measures of impulsive behavior, Neuropsychopharmacol. Off. Publ. Am. Coll. Neuropsychopharmacol. 29 (2004) 1331–1343. doi:10.1038/sj.npp.1300434.

[26] R.L. Barlow, J. Alsiö, B. Jupp, R. Rabinovich, S. Shrestha, A.C. Roberts, T.W. Robbins, J.W. Dalley, Markers of serotonergic function in the orbitofrontal cortex and dorsal raphé nucleus predict individual variation in spatial-discrimination serial reversal learning, Neuropsychopharmacol. Off. Publ. Am. Coll. Neuropsychopharmacol. 40 (2015) 1619–1630. doi:10.1038/npp.2014.335.

[27] F. Loiseau, A. Dekeyne, M.J. Millan, Pro-cognitive effects of 5-HT6 receptor antagonists in the social recognition procedure in rats: implication of the frontal cortex, Psychopharmacology (Berl.). 196 (2008) 93–104. doi:10.1007/s00213-007-0934-5.

[28] S.F. de Boer, D. Caramaschi, D. Natarajan, J.M. Koolhaas, The vicious cycle towards violence: focus on the negative feedback mechanisms of brain serotonin neurotransmission, Front. Behav. Neurosci. 3 (2009) 52. doi:10.3389/neuro.08.052.2009.

[29] L. Lewejohann, V. Kloke, R.S. Heiming, F. Jansen, S. Kaiser, A. Schmitt, K.P. Lesch, N. Sachser, Social status and day-to-day behaviour of male serotonin transporter knockout mice, Behav. Brain Res. 211 (2010) 220–228. doi:10.1016/j.bbr.2010.03.035.

[30] C.R. McKittrick, D.C. Blanchard, R.J. Blanchard, B.S. McEwen, R.R. Sakai, Serotonin receptor binding in a colony model of chronic social stress, Biol. Psychiatry. 37 (1995) 383–393.

[31] K. Kaplan, A.E. Echert, B. Massat, M.M. Puissant, O. Palygin, A.M. Geurts, M.R. Hodges, Chronic central serotonin depletion attenuates ventilation and body temperature in young but not adult Tph2 knockout rats, J. Appl. Physiol. Bethesda Md 1985. 120 (2016) 1070–1081. doi:10.1152/japplphysiol.01015.2015.

[32] J.P. Aggleton, The ability of different strains of rats to acquire a visual nonmatching-to-sample task, Psychobiology. 24 (1996) 44–48. doi:10.3758/BF03331952.

[33] A.H. Tuttle, V.M. Philip, E.J. Chesler, J.S. Mogil, Comparing phenotypic variation between inbred and outbred mice, Nat. Methods. 15 (2018) 994. doi:10.1038/s41592-018-0224-7.

[34] J. Myerson, L. Green, M. Warusawitharana, Area under the curve as a measure of discounting, J. Exp. Anal. Behav. 76 (2001) 235–243. doi:10.1901/jeab.2001.76-235.

[35] F. Zoratto, E. Sinclair, A. Manciocco, A. Vitale, G. Laviola, W. Adriani, Individual differences in gambling proneness among rats and common marmosets: an automated choice task, BioMed Res. Int. 2014 (2014) 927685. doi:10.1155/2014/927685.

[36] F. Dellu-Hagedorn, Relationship between impulsivity, hyperactivity and working memory: a differential analysis in the rat, Behav. Brain Funct. 2 (2006) 10. doi:10.1186/1744-9081-2-10.

[37] H. Arakawa, D.C. Blanchard, R.J. Blanchard, Colony formation of C57BL/6J mice in visible burrow system: Identification of eusocial behaviors in a background strain for genetic animal models of autism, Behav. Brain Res. 176 (2007) 27–39. doi:10.1016/j.bbr.2006.07.027.

[38] O. Burman, D. Owen, U. AbouIsmail, M. Mendl, Removing individual rats affects indicators of welfare in the remaining group members, Physiol. Behav. 93 (2008) 89–96. doi:10.1016/j.physbeh.2007.08.001.

[39] D.J. Rademacher, A.L. Schuyler, C.K. Kruschel, R.E. Steinpreis, Effects of cocaine and putative atypical antipsychotics on rat social behavior: An ethopharmacological study, Pharmacol. Biochem. Behav. 73 (2002) 769–778. doi:10.1016/S0091-3057(02)00904-8.

[40] Whishaw, Ian Q, Kolb Bryan, Behavior of the Laboratory Rat: A Handbook with Tests - Oxford Scholarship, (2004). http://www.oxfordscholarship.com/view/10.1093/acprof:oso/9780195162851.001.0001/acprof-9780195162851.

[41] M. Lepschy, C. Touma, R. Hruby, R. Palme, Non-invasive measurement of adrenocortical activity in male and female rats, Lab. Anim. 41 (2007) 372–387. doi:10.1258/002367707781282730.

[42] C. Touma, N. Sachser, E. Möstl, R. Palme, Effects of sex and time of day on metabolism and excretion of corticosterone in urine and feces of mice, Gen. Comp. Endocrinol. 130 (2003) 267–278.

[43] H. Shahar-Gold, R. Gur, S. Wagner, Rapid and Reversible Impairments of Short- and Long-Term Social Recognition Memory Are Caused by Acute Isolation of Adult Rats via Distinct Mechanisms, PLoS ONE. 8 (2013) e65085. doi:10.1371/journal.pone.0065085.

[44] R Core Team, R: The R Project for Statistical Computing, R Foundation for Statistical Computing, Vienna, Austria. https://www.R-project.org/, (2018). https://www.r-project.org/.

[45] Maxime Hervé, RVAideMemoire: Testing and Plotting Procedures for Biostatistics, (2018). https://CRAN.R-project.org/package=RVAideMemoire.

[46] B. Wheeler, M. Torchiano, lmPerm: Permutation Tests for Linear Models, (2016). https://CRAN.R-project.org/package=lmPerm.

[47] A. Liaw, M. Wiener, Classification and Regression by randomForest, (2002). https://CRAN.R-project.org/doc/Rnews/.

[48] N. Haines, J. Vassileva, W.-Y. Ahn, The Outcome-Representation Learning Model: A Novel Reinforcement Learning Model of the Iowa Gambling Task, Cogn. Sci. (2018). doi:10.1111/cogs.12688.

[49] M. Balconi, R. Finocchiaro, Y. Canavesio, Reward Sensitivity (Behavioral Activation System), Cognitive, and Metacognitive Control in Gambling Behavior: Evidences From Behavioral, Feedback-Related Negativity, and P300 Effect, J. Neuropsychiatry Clin. Neurosci. 27 (2015) 219–227. doi:10.1176/appi.neuropsych.14070165.

[50] A. Bechara, H. Damasio, Decision-making and addiction (part I): impaired activation of somatic states in substance dependent individuals when pondering decisions with negative future consequences, Neuropsychologia. 40 (2002) 1675–1689. doi:10.1016/S0028-3932(02)00015-5.

[51] M. Brand, E. Kalbe, K. Labudda, E. Fujiwara, J. Kessler, H.J. Markowitsch, Decision-making impairments in patients with pathological gambling, Psychiatry Res. 133 (2005) 91–99. doi:10.1016/j.psychres.2004.10.003.

[52] Y.-T. Kim, H. Sohn, J. Jeong, Delayed transition from ambiguous to risky decision making in alcohol dependence during Iowa Gambling Task, Psychiatry Res. 190 (2011) 297–303. doi:10.1016/j.psychres.2011.05.003.

[53] G. Fond, S. Bayard, D. Capdevielle, J. Del-Monte, N. Mimoun, A. Macgregor, J.-P. Boulenger, M.- C. Gely-Nargeot, S. Raffard, A further evaluation of decision-making under risk and under ambiguity in schizophrenia, Eur. Arch. Psychiatry Clin. Neurosci. 263 (2013) 249–257. doi:10.1007/s00406-012-0330-y.

[54] L. Zhang, J. Tang, Y. Dong, Y. Ji, R. Tao, Z. Liang, J. Chen, Y. Wu, K. Wang, Similarities and Differences in Decision-Making Impairments between Autism Spectrum Disorder and Schizophrenia, Front. Behav. Neurosci. 9 (2015) 259. doi:10.3389/fnbeh.2015.00259.

[55] H.W. Kim, J.I. Kang, K. Namkoong, K. Jhung, R.Y. Ha, S.J. Kim, Further evidence of a dissociation between decision-making under ambiguity and decision-making under risk in obsessive-compulsive disorder, J. Affect. Disord. 176 (2015) 118–124. doi:10.1016/j.jad.2015.01.060.

[56] N. Adjeroud, J. Besnard, C. Verny, A. Prundean, C. Scherer, B. Gohier, D. Bonneau, N.E. Massioui, P. Allain, Dissociation between decision-making under risk and decision-making under ambiguity in premanifest and manifest Huntington’s disease, Neuropsychologia. 103 (2017) 87–95. doi:10.1016/j.neuropsychologia.2017.07.011.

[57] E.A. Deisenhammer, S.K. Schmid, G. Kemmler, B. Moser, M. Delazer, Decision making under risk and under ambiguity in depressed suicide attempters, depressed non-attempters and healthy controls, J. Affect. Disord. 226 (2018) 261–266. doi:10.1016/j.jad.2017.10.012.

[58] J.E. Mekarski, Main effects of current and pimozide on prepared and learned self-stimulation behaviors are on performance not reward, Pharmacol. Biochem. Behav. 31 (1988) 845–853.

[59] J.E. Goeders, K.S. Murnane, M.L. Banks, W.E. Fantegrossi, Escalation of food-maintained responding and sensitivity to the locomotor stimulant effects of cocaine in mice, Pharmacol. Biochem. Behav. 93 (2009) 67–74. doi:10.1016/j.pbb.2009.04.008.

[60] R.J. Blanchard, L. Dulloog, C. Markham, O. Nishimura, J. Nikulina Compton, A. Jun, C. Han, D.C. Blanchard, Sexual and aggressive interactions in a visible burrow system with provisioned burrows, Physiol. Behav. 72 (2001) 245–254.

[61] J.F. Davis, E.G. Krause, S.J. Melhorn, R.R. Sakai, S.C. Benoit, Dominant rats are natural risk takers and display increased motivation for food reward, Neuroscience. 162 (2009) 23–30. doi:10.1016/j.neuroscience.2009.04.039.

[62] B. Buwalda, J.M. Koolhaas, S.F. de Boer, Trait aggressiveness does not predict social dominance of rats in the Visible Burrow System, Physiol. Behav. 178 (2017) 134–143. doi:10.1016/j.physbeh.2017.01.008.

[63] M.I. Cordero, C. Sandi, Stress Amplifies Memory for Social Hierarchy, Front. Neurosci. 1 (2007) 175–184. doi:10.3389/neuro.01.1.1.013.2007.

[64] B. Jupp, J.E. Murray, E.R. Jordan, J. Xia, M. Fluharty, S. Shrestha, T.W. Robbins, J.W. Dalley, Social dominance in rats: effects on cocaine self-administration, novelty reactivity and dopamine receptor binding and content in the striatum, Psychopharmacology (Berl.). 233 (2016) 579–589. doi:10.1007/s00213-015-4122-8.

[65] D.C. Blanchard, R.L. Spencer, S.M. Weiss, R.J. Blanchard, B. McEwen, R.R. Sakai, Visible burrow system as a model of chronic social stress: behavioral and neuroendocrine correlates, Psychoneuroendocrinology. 20 (1995) 117–134.

[66] M.J. Ramirez, Behavioral parameters of social dominance in rats, Bull. Psychon. Soc. 15 (1980) 96–98. doi:10.3758/BF03334477.

[67] N. So, B. Franks, S. Lim, J.P. Curley, A Social Network Approach Reveals Associations between Mouse Social Dominance and Brain Gene Expression, PLoS ONE. 10 (2015). doi:10.1371/journal.pone.0134509.

[68] N. Alia-Klein, R.Z. Goldstein, A. Kriplani, J. Logan, D. Tomasi, B. Williams, F. Telang, E. Shumay, A. Biegon, I.W. Craig, F. Henn, G.-J. Wang, N.D. Volkow, J.S. Fowler, Brain monoamine oxidase A activity predicts trait aggression, J. Neurosci. Off. J. Soc. Neurosci. 28 (2008) 5099–5104. doi:10.1523/JNEUROSCI.0925-08.2008.

[69] F. Jollant, C. Buresi, S. Guillaume, I. Jaussent, F. Bellivier, M. Leboyer, D. Castelnau, A. Malafosse, P. Courtet, The influence of four serotonin-related genes on decision-making in suicide attempters, Am. J. Med. Genet. Part B Neuropsychiatr. Genet. Off. Publ. Int. Soc. Psychiatr. Genet. 144B (2007) 615–624. doi:10.1002/ajmg.b.30467.

[70] E. Anderzhanova, T. Kirmeier, C.T. Wotjak, Animal models in psychiatric research: The RDoC system as a new framework for endophenotype-oriented translational neuroscience, Neurobiol. Stress. 7 (2017) 47–56. doi:10.1016/j.ynstr.2017.03.003.

[71] T. Insel, B. Cuthbert, M. Garvey, R. Heinssen, D.S. Pine, K. Quinn, C. Sanislow, P. Wang, Research domain criteria (RDoC): toward a new classification framework for research on mental disorders, Am. J. Psychiatry. 167 (2010) 748–751. doi:10.1176/appi.ajp.2010.09091379.

[72] M.F.W. Festing, Evidence Should Trump Intuition by Preferring Inbred Strains to Outbred Stocks in Preclinical Research, ILAR J. 55 (2014) 399–404. doi:10.1093/ilar/ilu036.

[73] A.D. Lê, H. Kalant, Intravenous self-administration of alcohol in rats-problems with translation to humans, Addict. Biol. 22 (2017) 1665–1681. doi:10.1111/adb.12429.

[74] M. Shoaib, R. Spanagel, T. Stohr, T.S. Shippenberg, Strain differences in the rewarding and dopamine-releasing effects of morphine in rats, Psychopharmacology (Berl.). 117 (1995) 240–247.

[75] A.M. Cano, E.S. Murphy, G. Lupfer, Delay discounting predicts binge-eating in Wistar rats, Behav. Processes. 132 (2016) 1–4. doi:10.1016/j.beproc.2016.08.011.

[76] U. Datta, M. Martini, M. Fan, W. Sun, Compulsive sucrose- and cocaine-seeking behaviors in male and female Wistar rats, Psychopharmacology (Berl.). 235 (2018) 2395–2405. doi:10.1007/s00213-018-4937-1.

[77] T. Brand, R. Spanagel, M. Schneider, Decreased reward sensitivity in rats from the Fischer344 strain compared to Wistar rats is paralleled by differences in endocannabinoid signaling, PloS One. 7 (2012) e31169. doi:10.1371/journal.pone.0031169.

[78] L. Brimberg, S. Flaisher-Grinberg, E.A. Schilman, D. Joel, Strain differences in “compulsive” lever-pressing, Behav. Brain Res. 179 (2007) 141–151. doi:10.1016/j.bbr.2007.01.014.

[79] I. Dela Peña, I.J. Dela Peña, J.B. de la Peña, H.J. Kim, C.Y. Shin, D.H. Han, B.-N. Kim, J.H. Ryu, J.H. Cheong, Methylphenidate and Atomoxetine-Responsive Prefrontal Cortical Genetic Overlaps in “Impulsive” SHR/NCrl and Wistar Rats, Behav. Genet. 47 (2017) 564–580. doi:10.1007/s10519-017-9861-3.

[80] M. Casarrubea, V. Roy, F. Sorbera, M.S. Magnusson, A. Santangelo, A. Arabo, G. Crescimanno, Significant divergences between the temporal structure of the behavior in Wistar and in the spontaneously more anxious DA/Han strain of rats tested in elevated plus maze, Behav. Brain Res. 250 (2013) 166–173. doi:10.1016/j.bbr.2013.05.016.

[81] A.O. Mechan, P.M. Moran, M. Elliott, A.J. Young, M.H. Joseph, R. Green, A comparison between Dark Agouti and Sprague-Dawley rats in their behaviour on the elevated plus-maze, open-field apparatus and activity meters, and their response to diazepam, Psychopharmacology (Berl.). 159 (2002) 188–195. doi:10.1007/s002130100902.

[82] American Psychiatric Association, Diagnostic and Statistical Manual of Mental Disorders, Fifth Edition. Washington DC, (2013). https://www.psychiatry.org/psychiatrists/practice/dsm.

