## Supplementary results for "Inter-Individual and Inter-Strain Differences in Cognitive and Social Abilities of Dark Agouti and Wistar Han Rats"

### Supplementary material

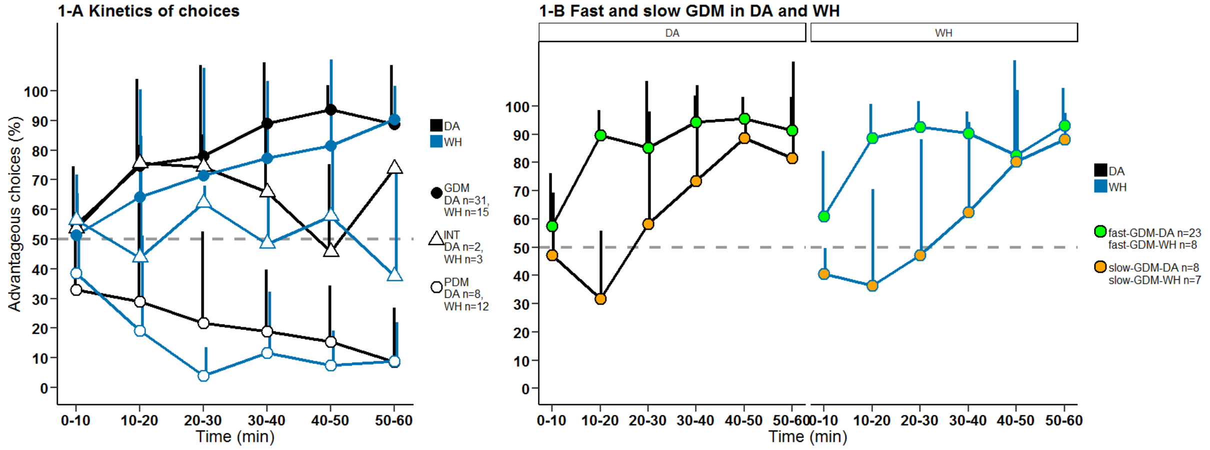


**Supplementary Figure 1. RGT A-** Kinetics of choices of GDMs, INTs and PDMs of the 2 strains. **B-** Fast and slow GDMs of the 2 strains.

The choices over time of the GDMs of both strains were comparable (Supplementary Figure 1A). However, while the DA GDMs and WH GDMs started at chance level (50%) they showed different evolutions of choices over time (Wilcoxon rank sum test, W = 9466.5, p = 0.0069). DA GDMs clearly preferred the advantageous options after 10 min of test (Figure 1C, Wilcoxon sign test, DA: CI [77.8, 96.0], p = 0.00010), while WH GDMs showed such preference only after 30 min of test (Wilcoxon sign test, WH: CI [76.9, 95.5], p = 0,0018). The dynamics of choices of the PDM rats were similar between strains. DA PDMs and WH PDMs both started the test at chance level, and their preferences for the disadvantageous options were significant after 40 min for the DA PDMs (Wilcoxon sign test, CI [0, 33.3], p = 0,015) and after 20 min for the WH PDMs (Wilcoxon sign test, CI [0, 0], p = 0,00048).

In both strains, “fast” and “slow” GDMs could be identified (Supplementary Figure 1B). In DA rats, the majority of the GDMs were of the “fast” type (76%, n = 23/30), significantly and consistently preferring the advantageous options after 10 min of test. In the WH half of the GDMs only were of the “fast” type (53%, n = 8/15).

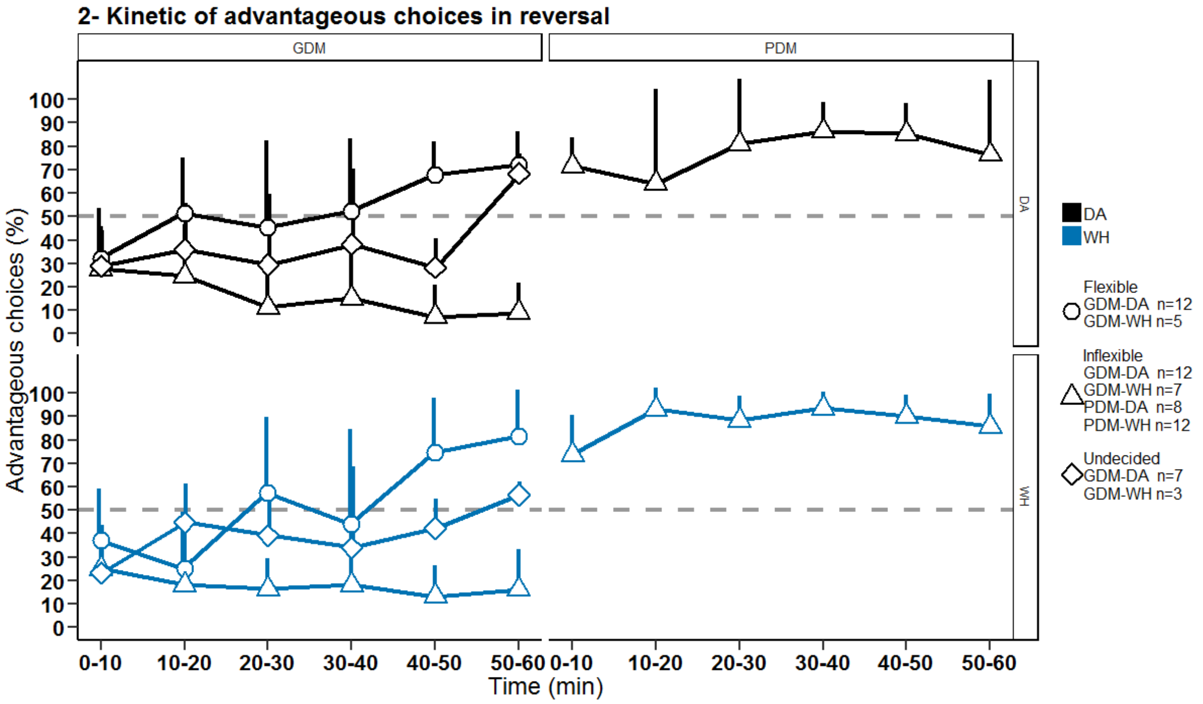


**Supplementary Figure 2. Kinetics of advantageous choices in the reversed-RGT.** The shape of the dot corresponds to the flexibility type: circle for flexible animals, triangle for inflexible animals and rhomb for undecided animals.

During the reversed-RGT, PDM rats kept choosing the hole(s) that they previously preferred during the RGT and despite the outcomes of these choices being the opposite to the outcomes experienced in the RGT (Supplementary Fig. 2). Flexible GDMs progressively (trial after trial) switched their spatial preference from the nose-poke holes previously associated with the advantageous options (in the RGT) to the nose-poke holes currently associated with the most advantageous options. Inflexible GDMs kept choosing the hole(s) previously preferred in the RGT although the outcomes of these options were not the ones preferred during the RGT. Undecided GDMs did not have a clear preference for each option at the end of the reversed-RGT.

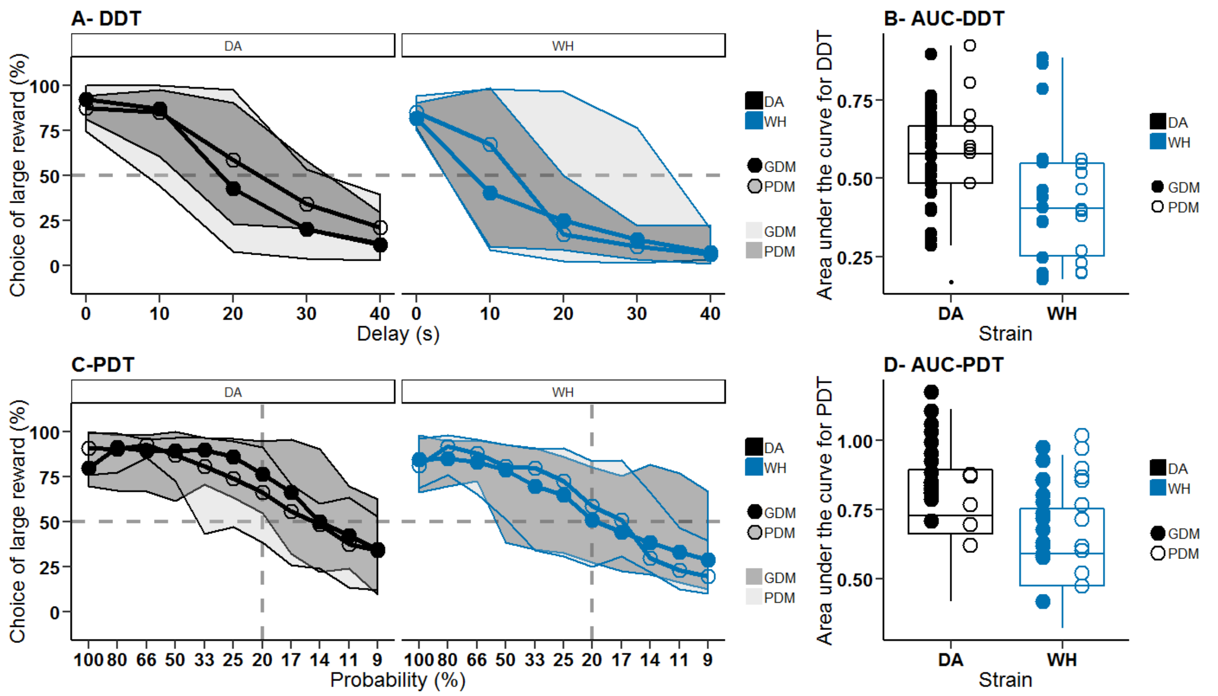


**Supplementary Figure 3. DDT A-**Kinetics of choices in the DDT. **B-**AUCs in the DDT. In the DDT, DA GDMs n = 30, DA PDMs n = 7, WH GDMs n = 15 and WH PDMs n = 12. **C-** Kinetics of choices in the PDT. **D-** AUCs in the PDT. In the PDT, DA GDMs n = 17, DA PDMs n = 5, WH GDMs n = 15 and WH PDMs n = 12. Filled dots represent the GDMs, empty dots represent the PDMs. The boxplots represent the entire population.

The tolerance to the delay was the same between GDM and PDM rats of the same strain. The DA rats switched preference at delay 20 s and the WH rats at delay 10 s (Supplementary Fig.3A). In both strains, PDMs and GDMs had similar AUCs (Supplementary Fig.3B). In the PDT, GDMs and PDMs of both strains behaved similarly toward the large-reward option at each probability-conditions (Supplementary Fig.3C). In both strains, PDMs and GDMs had similar AUCs (Supplementary Fig.3D).

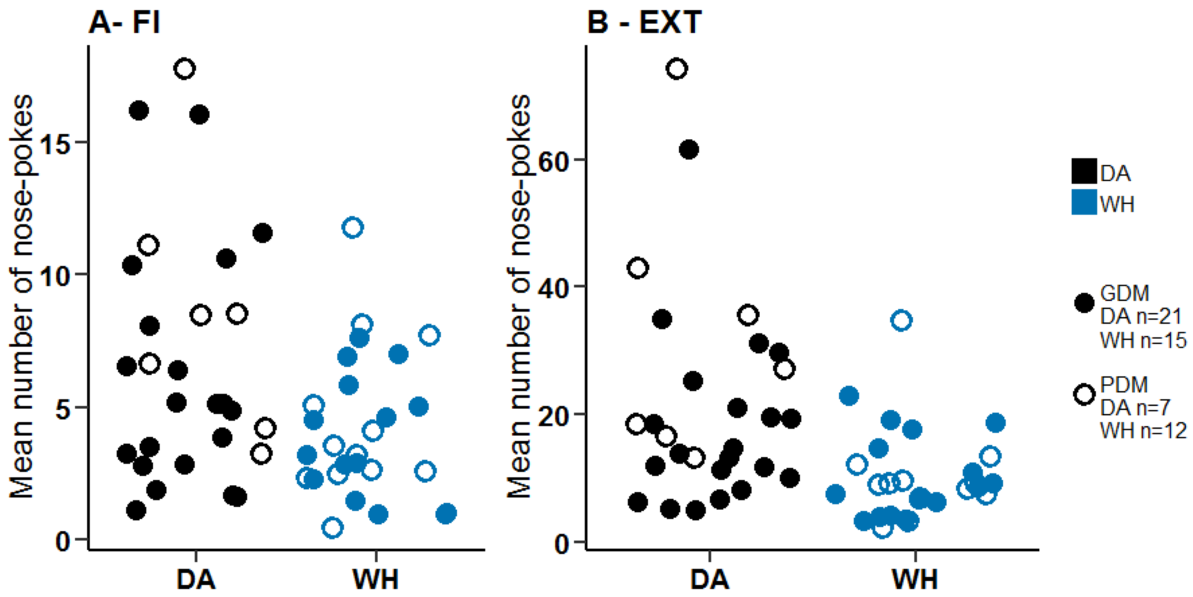


**Supplementary Figure 4. FI-EXT A-**Mean number of nose-pokes in the FI. **B-** Mean number of nose-pokes in the EXT. Filled dots represent the GDMs, empty dots represent the PDMs.

During FI, the mean number of nose pokes was equivalent between PDMs and GDMs of both strains (Supplementary Fig. 4A). However, during EXT, DA PDMs nose poked significantly more often than DA GDMs (Supplementary Fig. 4B, Wilcoxon rank sum test with continuity correction, $W = 35$, $p = 0.043$) which was not observed in WH.

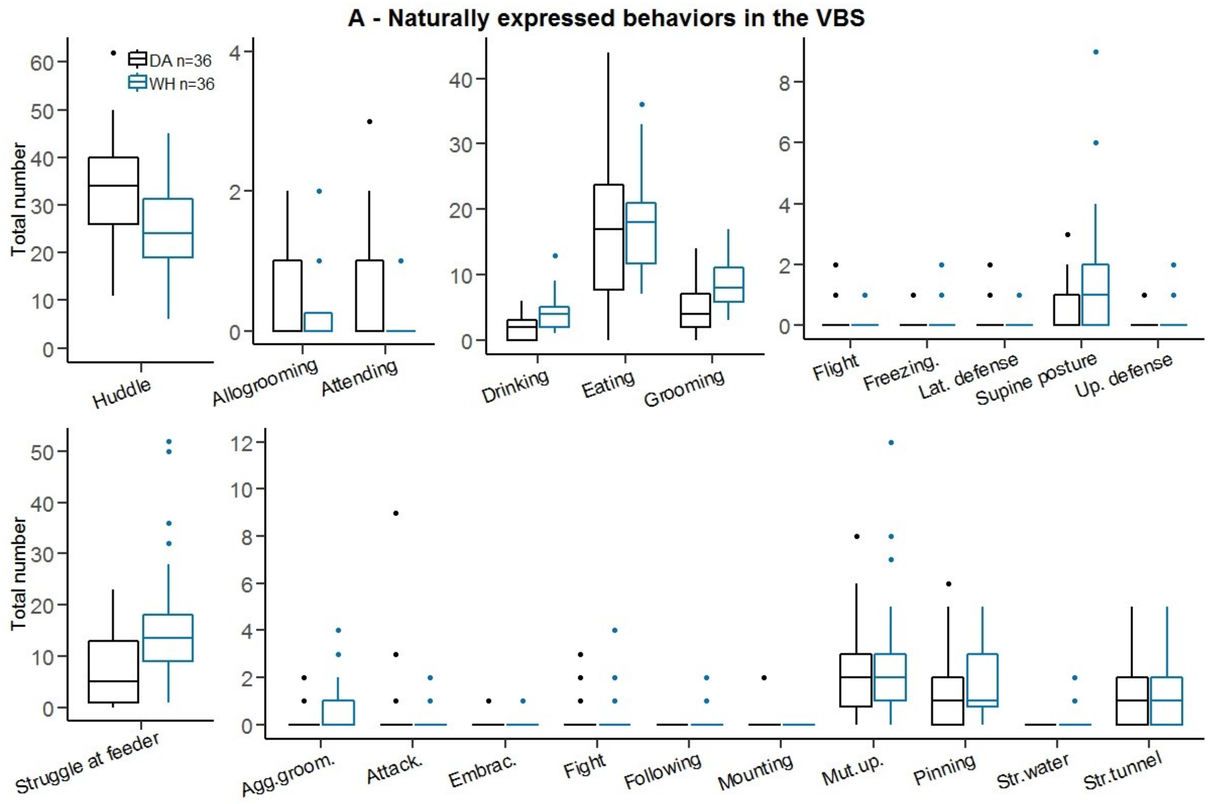


**Supplementary Figure 5. A**-Total number of occurrences of all scored behaviours in the VBS in DA and WH.

DA rats presented a higher number of affiliative behaviours than WH (data not shown, Wilcoxon rank sum test with continuity correction, W = 992, p = 0.00010), with a higher huddle behaviours (Supplementary Fig. 5A, Wilcoxon rank sum test with continuity correction, huddle: W = 984, p = 0.000156) and an equal number of allogrooming and attending than WH.

In average the number of maintenance behaviours was different between DA and WH rats (data not shown, Wilcoxon rank sum test with continuity correction, W = 445.5, p = 0.022), with the same number of eating bouts but a higher number of drinking and grooming behaviours in DA than in WH (Wilcoxon rank sum test with continuity correction, drinking: W = 264, p = 1.24e-05 and grooming: W = 287.5, p = 4.75e-05; Supplementary Fig.5A).

DA and WH rats expressed few defensive behaviours (Supplementary Fig.5A). DA rats were less aggressive than WH (data not shown, Wilcoxon rank sum test with continuity correction, 327, p = 0.0003005), with a lower number of struggle at feeder than WH rats (Supplementary

Fig.5A, struggle at feeder: Wilcoxon rank sum test with continuity correction, W= 313.5, p = 0.0001643). The number of other aggressive behaviours (i.e. aggressive grooming, attack, mutual upright posture, pinning and struggle in tunnel) were not different between the strains. The following behaviours (total occurrences in parentheses): mounting (2), embracing (5), following (8), struggle at water (10), lateral defense (7), freezing (8) upright defense (6) and flight (13) were very rarely observed in the VBS.

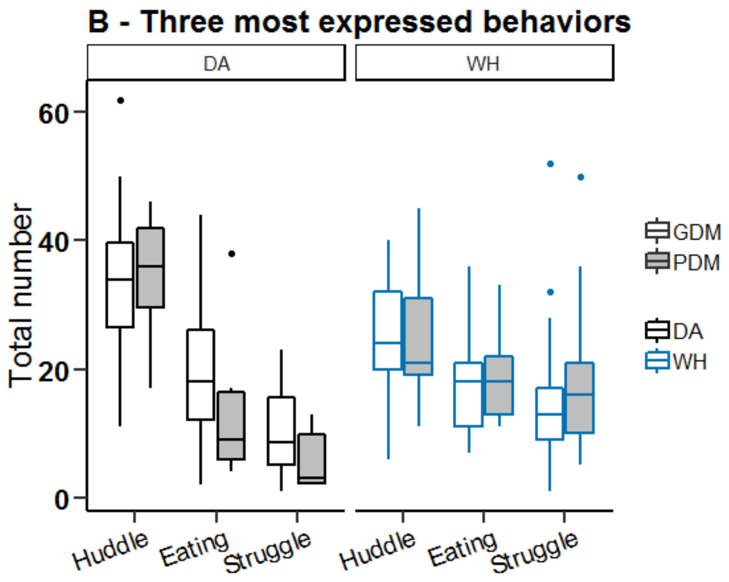


**Supplementary Figure 5. B**-Total number of occurrences of the three most observed behaviours in the Visible Burrow System.

The three most expressed behaviours were huddle, eating and struggle at feeder, and at similar rate in PDMs and GDMs of both strains (Supplementary Fig.5B).

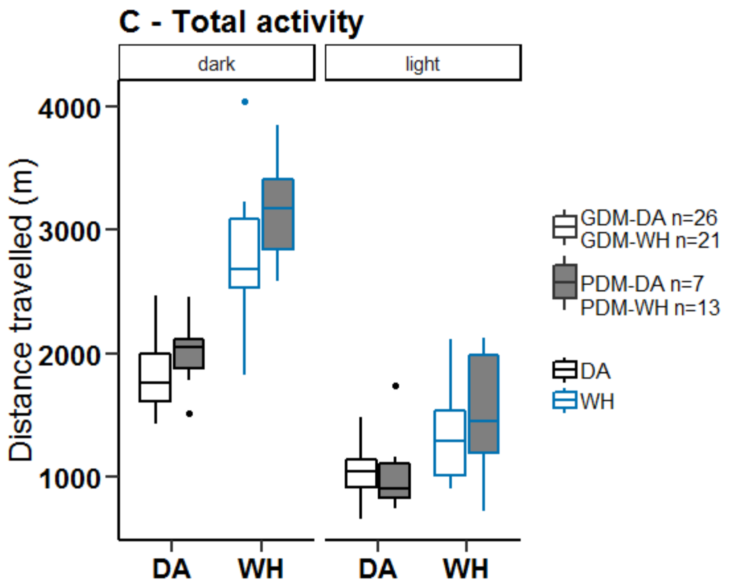


**Supplementary Figure 5. C**-Activity in the Visible Burrow System during dark and light phases.

The WH PDMs were more active than the WH GDMs during the dark phase (Wilcoxon rank sum test, W = 60, p = 0.0058), which was not the case during the light phase, or for the DA rats in both phases (Supplementary Fig.5C).

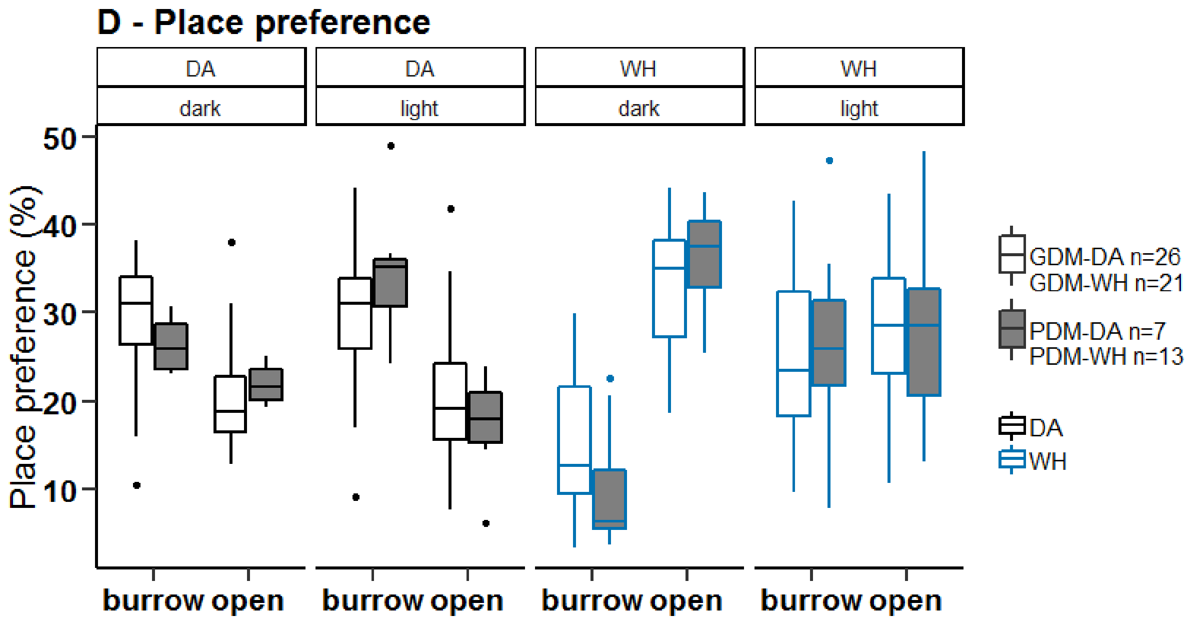


**Supplementary Figure 5. D**-Place preference in the Visible Burrow System during the dark and light phase.

The WH PDMs spent less time in the burrow during the dark phase than the WH GDMs (Wilcoxon rank sum test, W = 195, p = 0.038, Supplementary Fig.5D). This was only a tendency in DA PDMs (Wilcoxon rank sum test, W = 134, p-value = 0.060).

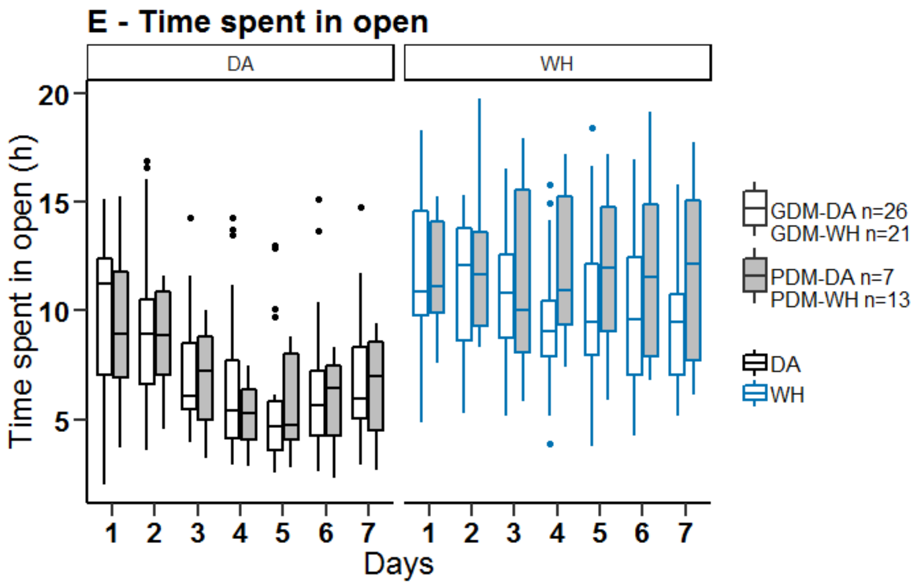


**Supplementary Figure 5. E-** Time spent in the open area per day in the VBS.

On day 4, WH PDMs spent significantly more time in the open area of the VBS than WH GDMs (non-parametric ANOVA with permutations, day 4, $p = 0.023$, Supplementary Fig.5F). It was not the case in DA.

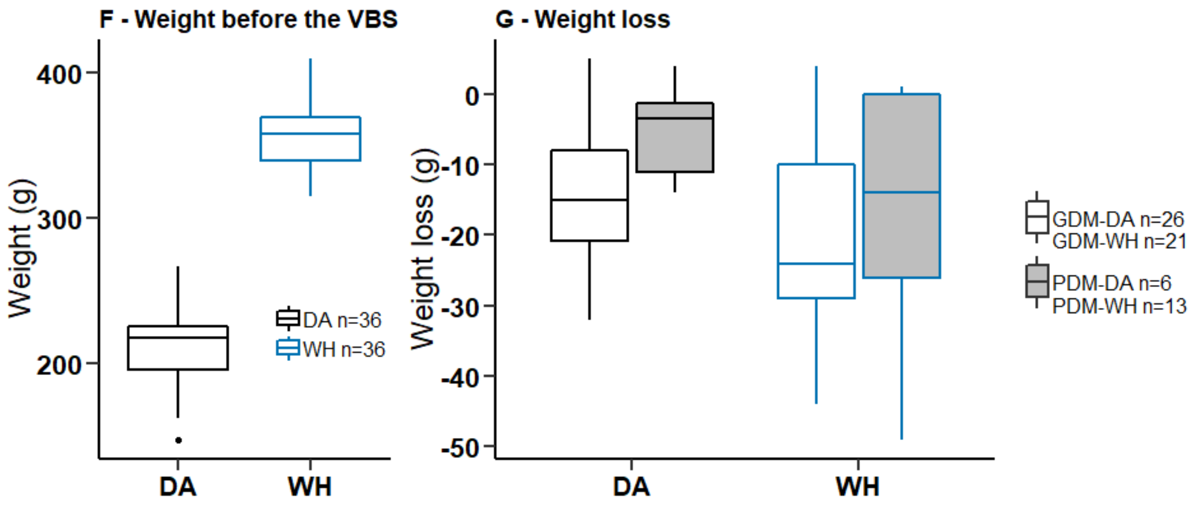


**Supplementary Figure 5. F-**Animals' weight before the VBS housing. **G-**Weight loss after the VBS housing.

DA and WH rats presented different weights before VBS housing, which was inherent to their strain (Supplementary Fig.5F). DA PDMs lost less weight than DA GDMs during the VBS (Wilcoxon rank sum test with continuity correction data, $W = 35$, $p = 0.039$), it was not the case in WH (Supplementary Fig.5G).

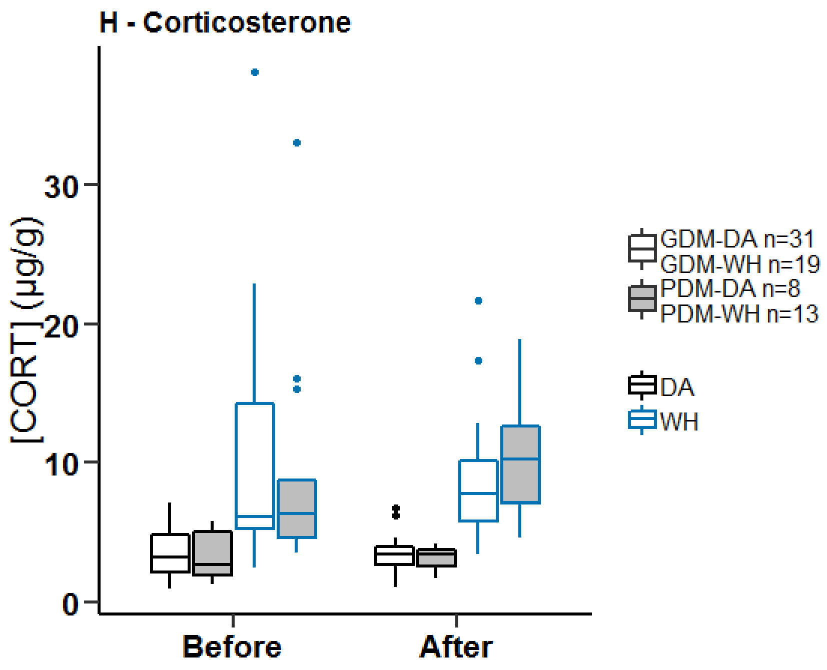


**Supplementary Figure 5. H**-Levels of corticosterone before and after VBS housing of PDMs and GDMs.

There were no differences in the levels of corticosterone between PDMs and GDMs of both strains either before or after the VBS housing.

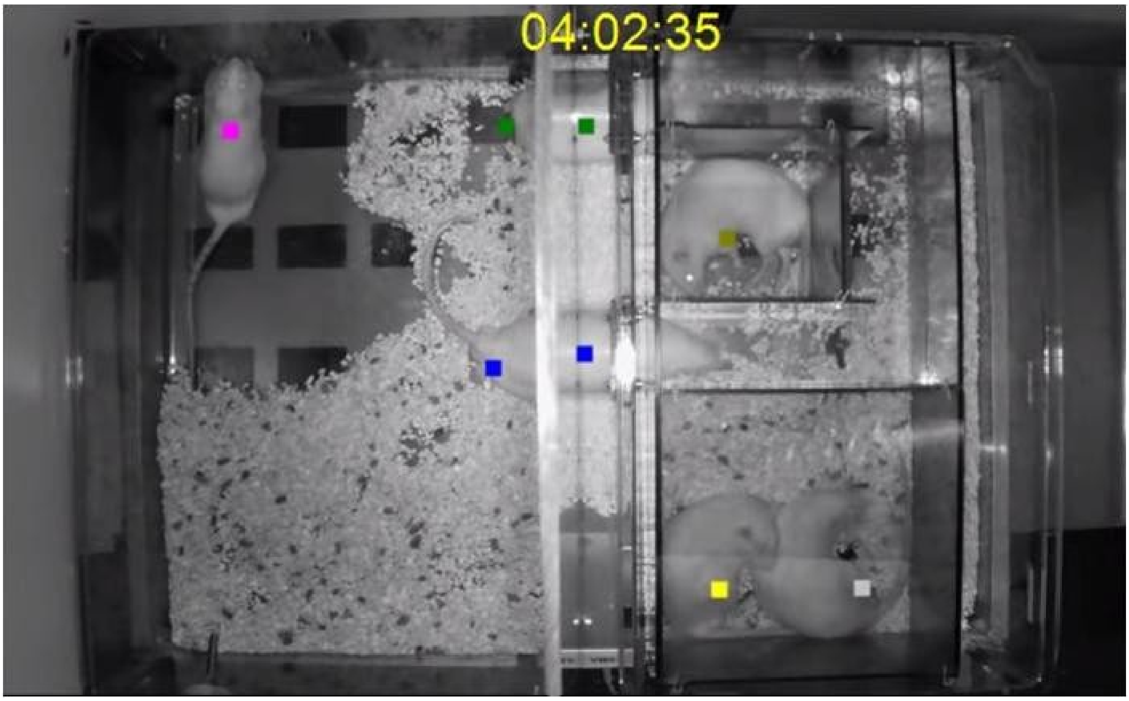


**Supplementary Figure 5. I**-Screenshot of the Visible Burrow System as seen in the recorded videos. Open area on the left and burrow system on the right. Color spots represent individual detections by the RFID reader.

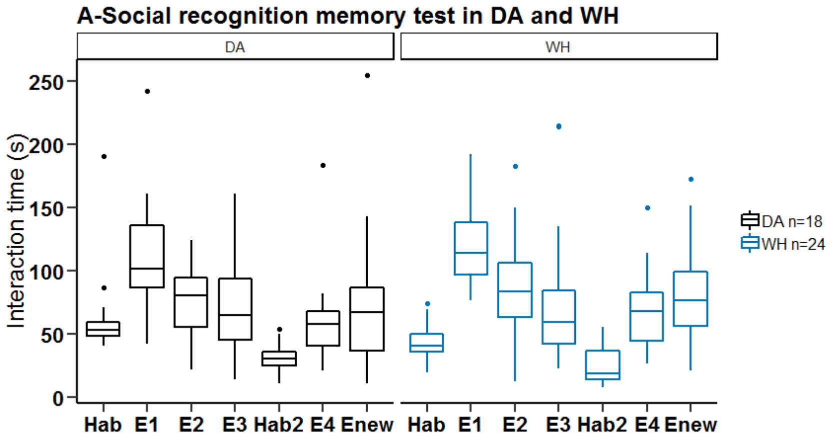


**B- Social Recognition setup**

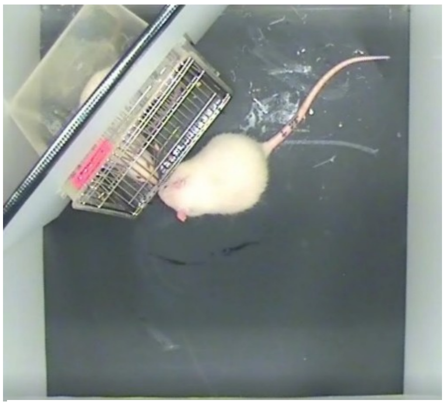

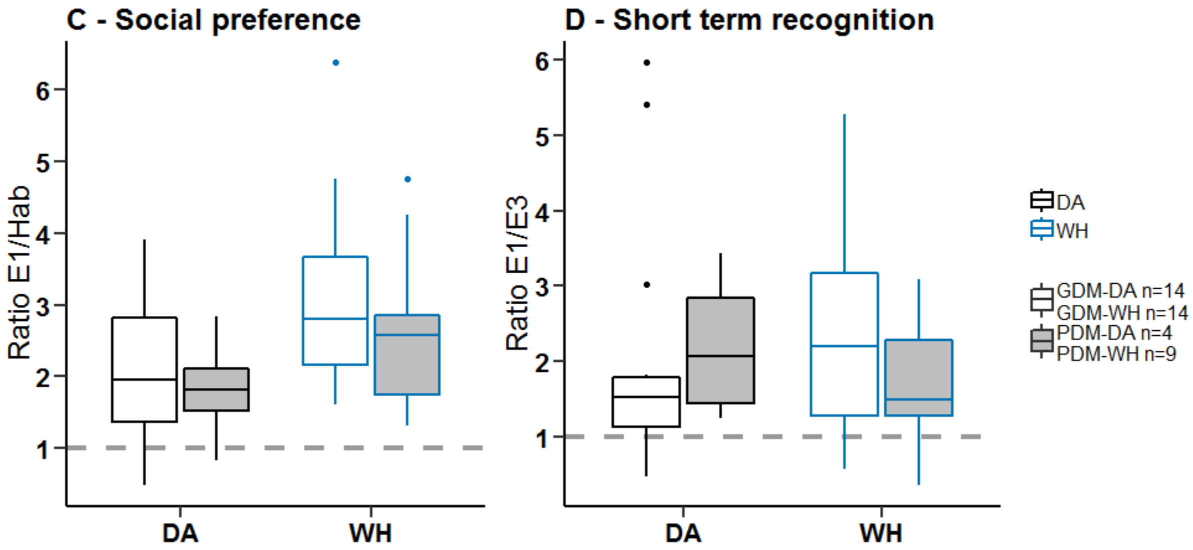


**Supplementary Figure 6. SRt A**-Interaction times in the SRt for all encounters. Hab: habituation phase and non-social cue (empty box), E1: first encounter with intruder1 (unfamiliar), E2: second encounter with intruder1 (familiar), E3: third encounter with intruder1 (familiar). Hab2: habituation phase, day 2 and non-social cue (empty box), E4: fourth encounter with intruder1 happening on day2 (familiar), Enew: first encounter with intruder2 on day2 (unfamiliar). **B**-Picture of the Social Recognition setup. **C**-Social preference of GDMs and PDMs represented as the ratio of exploration times in encounter 1 (E1) and habituation (Hab). **D**-Short term social recognition of GDMs and PDMs represented as the ratio of exploration times in encounter 1 (E1) and in encounter 3 (E3).

The presence of the intruder triggered an increase of the interaction time in E1 (social preference) compared to Hab. With the repetition of the encounters (E2, E3) with the same intruder the interaction time decreased indicating social recognition of the intruder by the subject (Supplementary Fig. 6A). Although we did not observe an increase in interaction time in Enew compared to E4, it is most likely that both strains were able to detect the unfamiliarity of the new intruder, but that the conditions of test in our set-up were not favourable to this measure (Supplementary Fig. 6A). On the picture of the setup (Supplementary Fig. 6B), the subject can be seen in the open field and the intruder can be seen in the cage. In the original study [42], the encounters were done in a cage in dim light with a free juvenile intruder, higher level of contact and investigation was allowed than in our setup. In both strains, the social preference ratio and short term memory ratio were not different in GDMs and PDMs (Supplementary Fig. 6C and D). Note that we had only four DA PDMs tested.

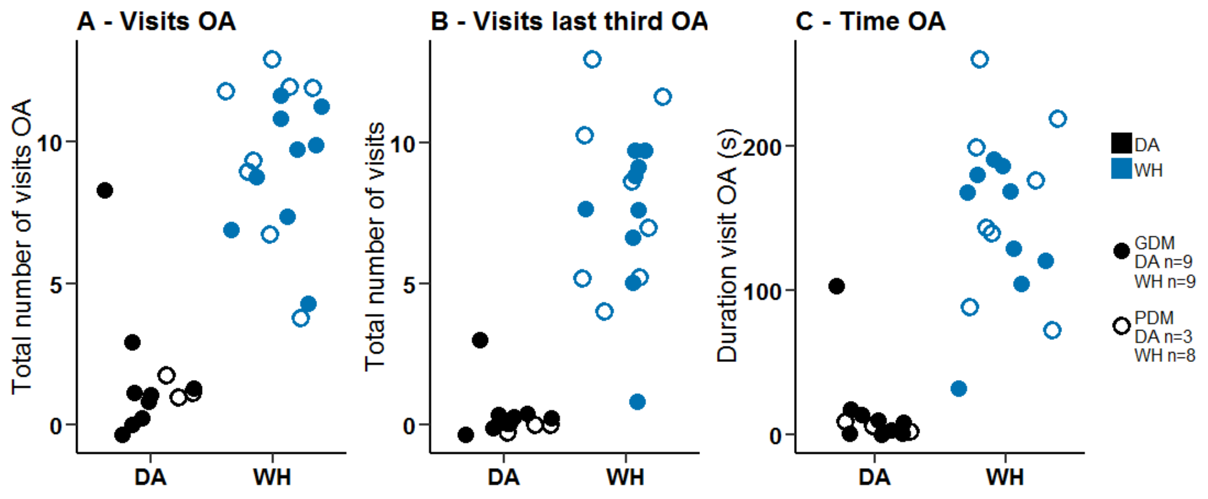


**Supplementary Figure 7. EPM A**-Total number of visits of the open arms. **B**-Total number of visits to the last third of the open arms. **C**-Total time spent in the open arms.

We did not observe differences between PDMs and GDMs in the parameters assessed in the EPM (Supplementary Fig. 7).

**Supplementary Table 1**: Eigenvalues of the principal component analysis

| Dimension | Eigenvalue | Variance (%) | Cumulative variance (%) |
|---|---|---|---|
| Dim 1 | 4,5 | 26,4 | 26,4 |
| Dim 2 | 2,4 | 14,3 | 40,7 |
| Dim 3 | 1,8 | 10,5 | 51,2 |
| Dim 4 | 1,4 | 8,5 | 59,7 |
| Dim 5 | 1,3 | 7,5 | 67,2 |
| Dim 6 | 1,2 | 7,1 | 74,3 |
| Dim 7 | 0,9 | 5,3 | 79,6 |
| Dim 8 | 0,7 | 3,8 | 83,5 |
| Dim 9 | 0,5 | 3,2 | 86,7 |
| Dim 10 | 0,5 | 2,7 | 89,4 |
| Dim 11 | 0,4 | 2,4 | 91,8 |
| Dim 12 | 0,4 | 2,1 | 93,9 |
| Dim 13 | 0,3 | 1,9 | 95,8 |
| Dim 14 | 0,2 | 1,5 | 97,2 |
| Dim 15 | 0,2 | 1,2 | 98,4 |
| Dim 16 | 0,2 | 1,0 | 99,4 |
| Dim 17 | 0,1 | 0,6 | 100,0 |

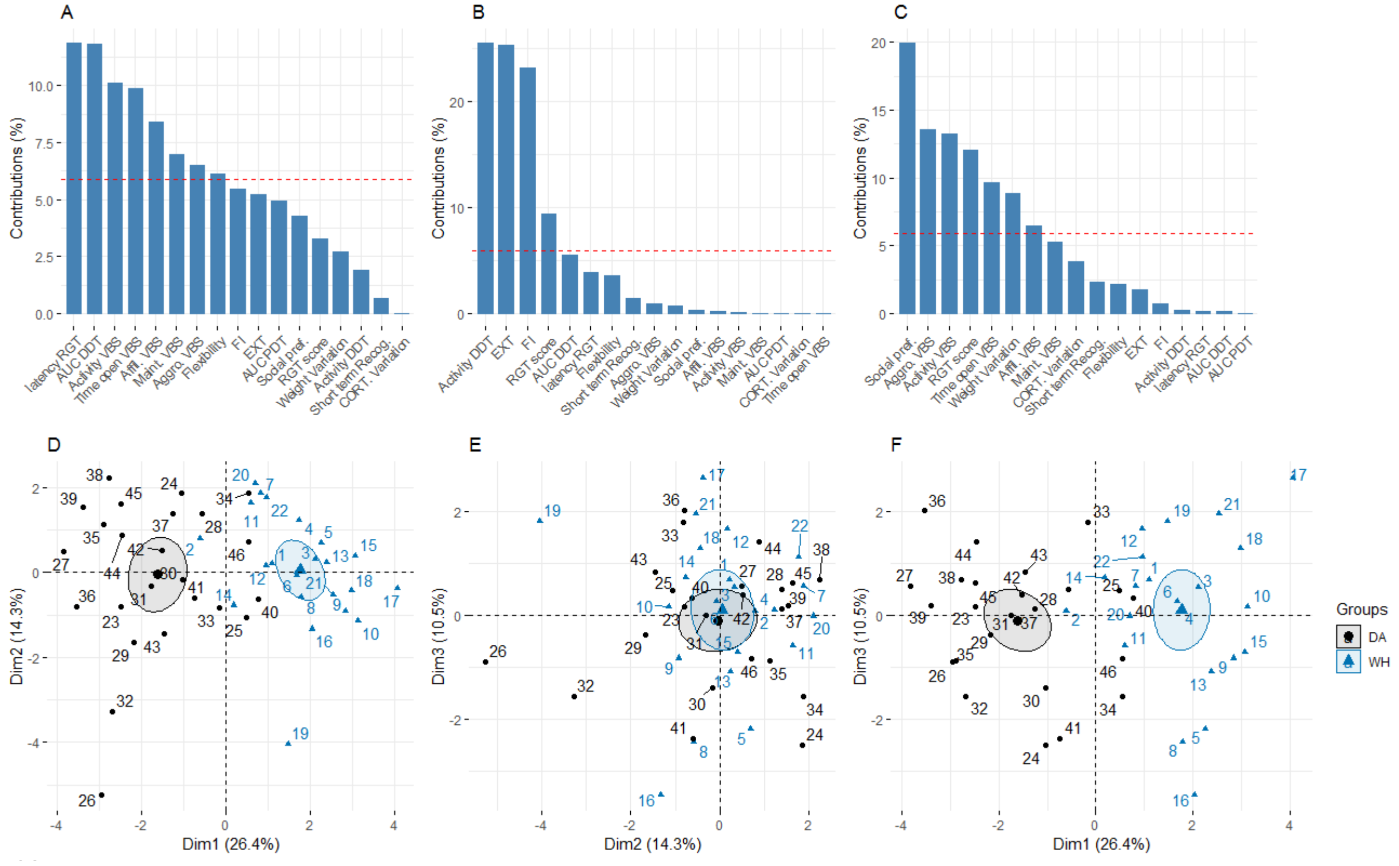


**Supplementary Figure 8.Principal Component Analysis.** Contribution of the variables to dimension 1 **(A)**, dimension 2 **(B)** and dimension 3 **(C)**. The red dashed line indicates the expected average contribution. Graph of the individual scores plotted along dimension 1 and dimension 2 **(D)**, dimension 2 and dimension 3 **(E)** and dimension 1 and dimension 3 **(F)**.

A principal component analysis (function prcomp) [43] was done on the same dataset as the one used for the Random Forest analysis (see main text; 2.4. Statistical analysis). Eight dimensions explained 80% of the variance of the dataset (see Supplementary table 1). The first dimension which explained 26% of the variance of the dataset, received the highest contribution from the latency to collect a reward in the RGT, the AUC of the DDT, the distance travelled in the VBS and the total time spent in the open area in the VBS (Supplementary Fig. 8A). These variables were identified with the Random forest classifier as the most important variables to discriminate between DA and WH rats (Fig. 4A). Indeed, the individuals of the DA and WH strains separated along the dimension 1 (Wilcoxon rank sum test, $W = 12$, $p < 0.001$, Supplementary Fig. 8D and F). The dimension 2 and 3 received high contributions from variables of lesser importance to discriminate between DA and WH rats (Supplementary Fig. 8B and C) and they did not allow to separate DA and WH rats on their respective dimensions (Supplementary Fig. 8E).
