## Supplementary results for "Inter-Individual and Inter-Strain Differences in Cognitive and Social Abilities of Dark Agouti and Wistar Han Rats"

SUPPLEMENT: Automated Visual Burrow System

**Fig.** Automated Visual Burrow System. (A) Dual-cage system consisting of the burrow cage (left, EU Type IV rat cage, Tecniplast Model 1354G) and home cage (right, Tecniplast 2000P). (B) 32-array of ID-chip sensors (PhenoSys, model RFID-long range). Sensors connected to a 32-channel RFID-controller (Phenosys, not shown) connected to a PC via an Ethernet cable. (C) Components of the burrow made from Perspex black 962, thickness 4 mm, except for the two long side walls with thickness 5 mm (parts 2 and 4).

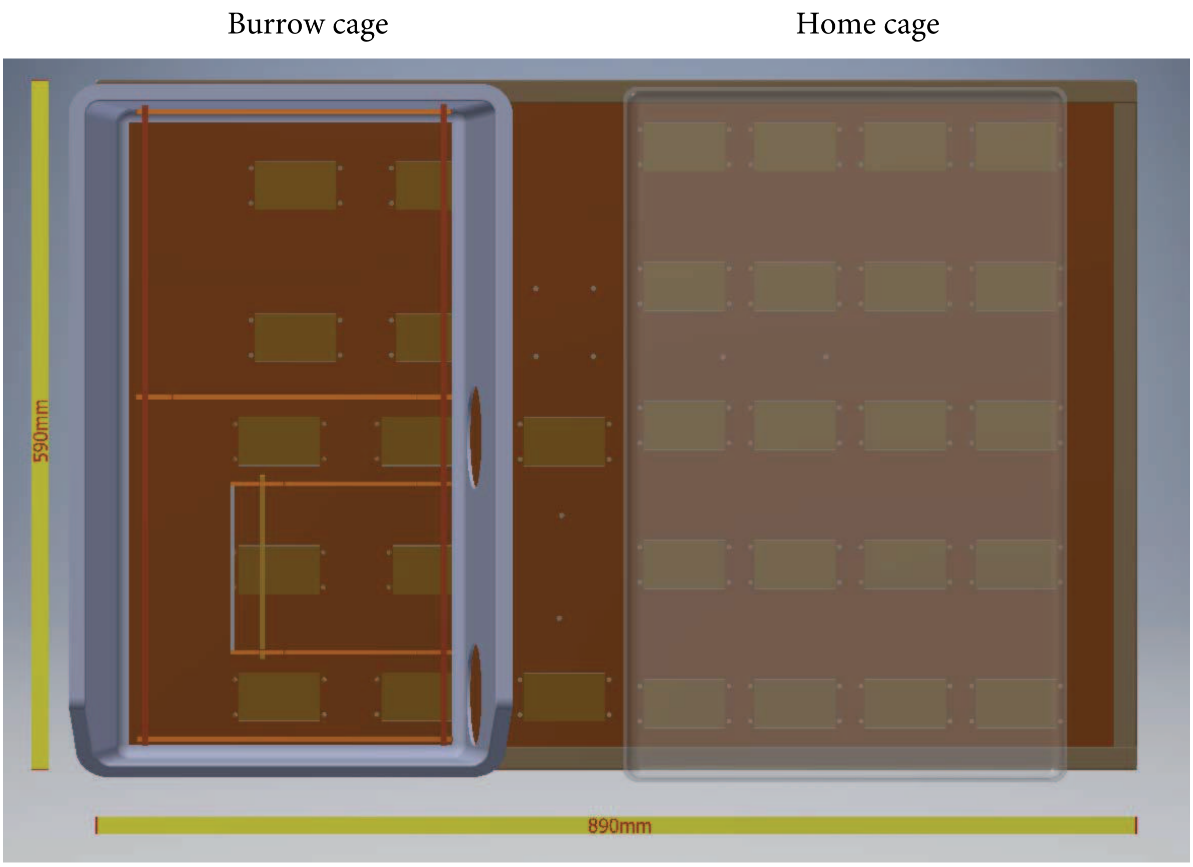


A: dual cage system

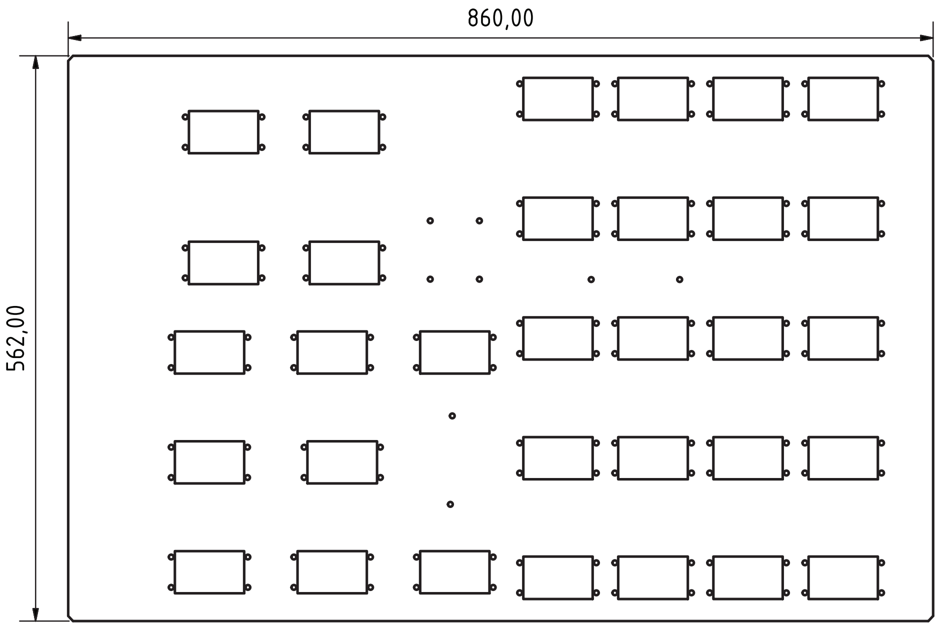


B: ID-chip reader plate

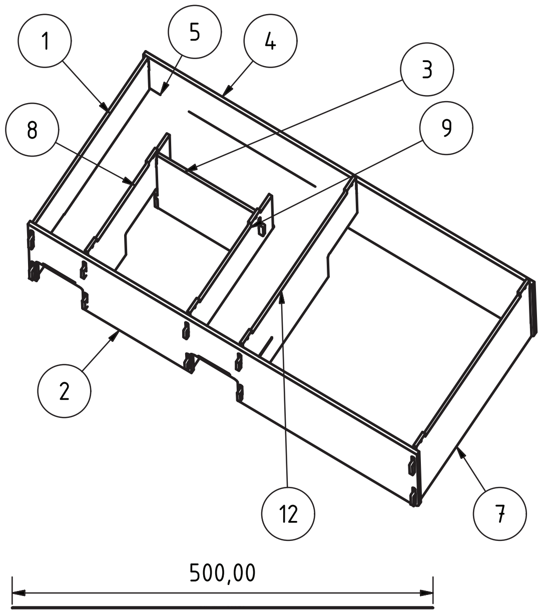


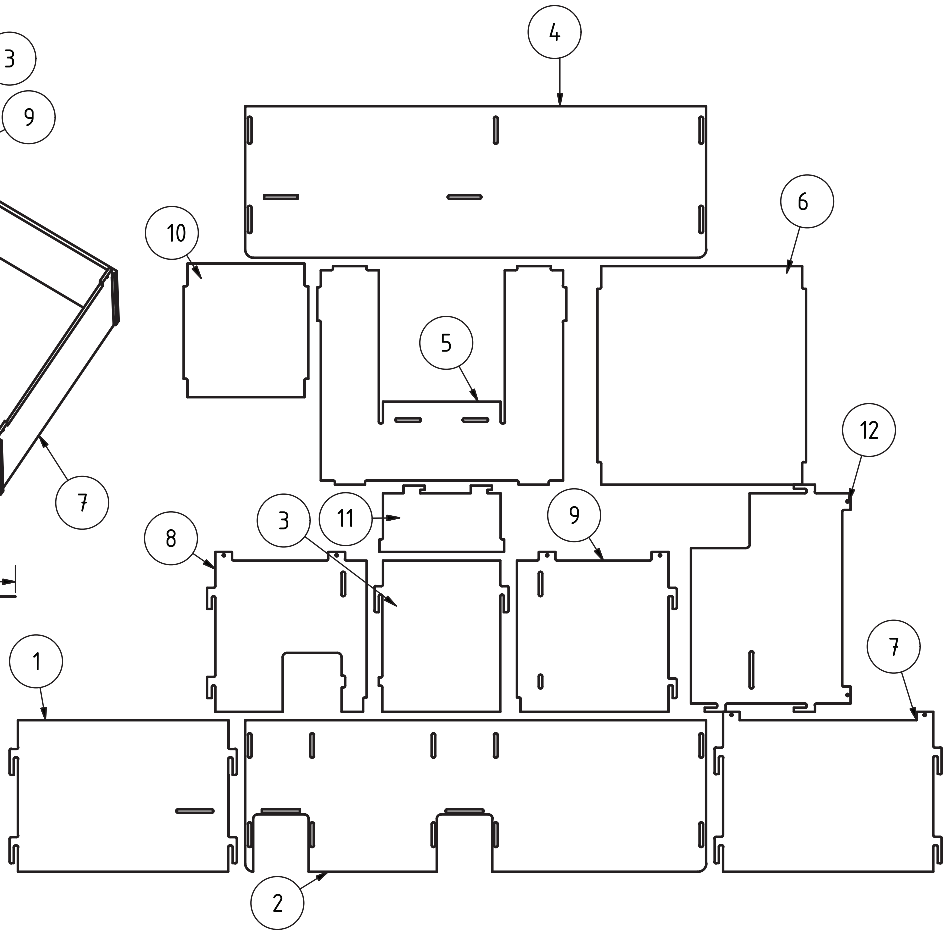


C: burrow components

This work is licensed under a [Creative Commons Attribution 4.0 International License](https://creativecommons.org/licenses/by/4.0/)
